# Circadian regulation of Homologous Recombination by Cryptochrome1-mediated dampening of DNA end resection

**DOI:** 10.1101/2023.01.22.524239

**Authors:** Amador Romero-Franco, Cintia Checa-Rodríguez, Sonia Jimeno, Maikel Castellano-Pozo, Paula Aguilera, Hector Miras, Amadeo Wals, Silvia Jimeno-González, Andres Joaquin Lopez-Contreras, Pablo Huertas

## Abstract

Genomic stability maintenance requires the repair of DNA breaks in the most accurate fashion. So, an exquisite regulatory network controls the choice between different repair mechanisms to maximize genome integrity. This relies mostly at the level of DNA end resection, the initial steps of the homologous recombination. On the other hand, numerous cellular activities follow a 24 h oscillation known as the circadian cycle. Thus, we explored the regulation of the choice between different DNA break repair pathways along the circadian cycle. Here we show that in human cells DNA resection shows a circadian oscillation, with a peak at early morning followed by a partial and progressive reduction until late afternoon. Such regulation depends on the circadian clock core component CRY1, which modulates the anti-resection activity of CCAR2 to limit CtIP at nightfall. Additionally, such regulation requires DNA-PK-mediated phosphorylation of CRY1. Finally, this circadian regulation impacts cancer progression and response to radiation therapy of specific tumours.

## INTRODUCTION

DNA double-strand break (DSB) repair is a critical aspect of cellular biology that has a great impact in different human diseases such as cancer ^1^. DSBs can be channelled through different repair pathways, with distinctive outcomes. Such pathways belong, mostly, to two different families: homology-driven repair (HDR), that exploits the presence of intact copies of the broken sequences as a template; and non-homologous end-joining (NHEJ), the simple ligation of two DNA ends with little or no processing ^2–4^. Both can result in either faithful or mistaken repair, thus the choice between them is tightly regulated. The best characterized regulatory step for DSB repair pathway choice is the activation of DNA end resection, a degradation of one strand of the broken DNA to create long tails of 3’ protruding ssDNA ^5,6^. Resection regulation relies mostly on the control of the core resection factor CtIP trough specific interactions and/or post-translational modifications ^7,8^. Once CtIP is activated, resection ensues and HDR prevails. If CtIP is blocked, resection is hampered and HDR is blocked. CCAR2 functions as a CtIP antagonist in resection regulation ^9^, acting as a downstream effector of the Shieldin complex ^10^. Both proteins interact constitutively, and only when CCAR2 is evicted from damaged chromatin, CtIP is unleashed, thus favouring resection ^9^.

Circadian rhythms are a series of periodic oscillations that follow the day/night cycle ^11^. Its presence is conserved throughout the whole photosensitive spectrum of living organisms ^12^. The circadian clock works as a built-in 24-hour oscillator that acts in a two-tier fashion: an intrinsic cellular circadian circuitry with a roughly 24 h period that can be synchronized in response to the external light, a process known as photoentrainment ^11^. In mammals, at the molecular level, the transcription factors BMAL1 and CLOCK stimulate the expression of the genes that code for the Period (PER1-3) and Cryptochrome (CRY1-2) families during the activation phase that, in humans, coincides with daylight hours. Then, PER and CRY heterodimers translocate into the nucleus where CRY negatively affect the transcriptional activity of BMAL1/CLOCK, mainly during night hours ^13^. Thus, a recurrent oscillation is created by sequential transcriptional activation/repression waves. Additional accessory loops have been characterized in the circadian clock, but the heterodimers BMAL1/CLOCK and PER/CRY build the core cellular oscillator ^14^. Importantly, the robustness of the circadian cycle varies among tissues, but it is also affected by the age of the individuals and even shows sex-dimorphic characteristics ^15^.

Here we show that DNA end resection and HDR, are regulated in a circadian fashion. Such regulation relies specifically and directly on a negative effect of the circadian factor CRY1. Indeed, resection activity peaks when CRY1 levels are low, corresponding in ideal circadian cycle in humans to early morning. Then it slowly decreases as CRY1 levels increase until a point that would correspond in humans to the evening. Finally, this is followed by an increase during what would correspond to the night in a correlation with a decline in CRY1 levels. Mechanistically, CRY1 is recruited to sites of DSBs and reinforces the inhibitory effect of CCAR2 over CtIP by potentiating CCAR2 retention on damaged chromatin. Furthermore, DNA-PK finetunes this circadian effect. So, when CRY1 is recruited to damaged chromatin is phosphorylated by DNA-PK, increasing its retention at DSBs. This locks CCAR2 there, curbing CtIP activity. Finally, this fine regulation of end resection affects the ability of cells to deal with DNA damaging agents in cellular models and mice xenografts, but more importantly in cancer clinical setups.

## RESULTS

### DNA end resection shows a circadian synchrony

As mentioned, all animal cells have a built-in circadian oscillator. Although *in vitro* grown cell cultures cannot be photoentrained, the addition of different drugs can reset them ^16,17^. We synchronized U2OS cells with either Dexamethasone (DEX) or Forskolin (FSK) for two hours, resetting the internal clock to the equivalent of “early morning” (Fig.1A), i.e when CRY levels are lower. Then, we irradiated them to induce DSBs at fixed time points after release, that represent different times of the day/different CRY1 levels (Fig. 1A). A control treated with the vehicle, i.e. an asynchronous population, was prepared in parallel. Circadian synchronization was followed by the levels of CRY1 and CRY2, that peaked between 8h (CRY2) or 12h (CRY1) after release (Fig. 1B for DEX). Indeed, such peak was due to circadian synchronization, as it was absent in the control experiment treated with the vehicle EtOH alone (Fig. 1C). Then, we analysed DNA end resection by checking the formation of RPA foci by immunofluorescence upon 10 Gy of irradiation. U2OS cells synchronized with DEX (blue line) showed a marked oscillation in resection proficiency, clearly distinct to the observed in the control treated with the vehicle EtOH (red line Fig. 1D; see Suppl. Fig. 1A for representative images). Whereas 6 and 18 hours after release the number of cells actively resecting was like the control, a marked reduction was readily observed at the 12 hours timepoint. Moreover, just after release resection was slightly elevated when compared with the asynchronous culture. Similar results were observed by adding FSK when compared with the mock treatment (Fig. 1E, Suppl. Fig. 1B), reinforcing the idea that was due to circadian synchronization and not an off-target effect of the drugs. It has been described that during the first 24h after release from DEX there are some effects that might be mediated by immediate-early/early gene expression and not due to a true circadian effect. To clarify this issue, we repeated the RPA foci formation experiment upon DEX synchronization and release at different time points during 48 h. As shown in Fig. 1F (see Suppl. Fig. 1C for representative images and Suppl. Table 1 for statistical analysis), the same pattern was observed both during the first and second days upon release, strengthening the idea that resection is more proficient during the times that correspond to the beginning of the circadian cycle, i.e. when CRY levels are low, and reduced at the time that correspond to CRY peaks. As we have never observed any oscillation of the foci in EtOH treated cells collected at different timepoints, from this point onward, only a single asynchronous sample was taken as a control and represented as a dashed line. Indeed, as expected, resection patterns behave opposite to the accumulation of the circadian protein CRY1, measured by immunofluorescence (Fig. 1G, Suppl. Table 1, Suppl. Fig. 1D) or western blot (Suppl. Fig. 1E).

**Figure 1.**
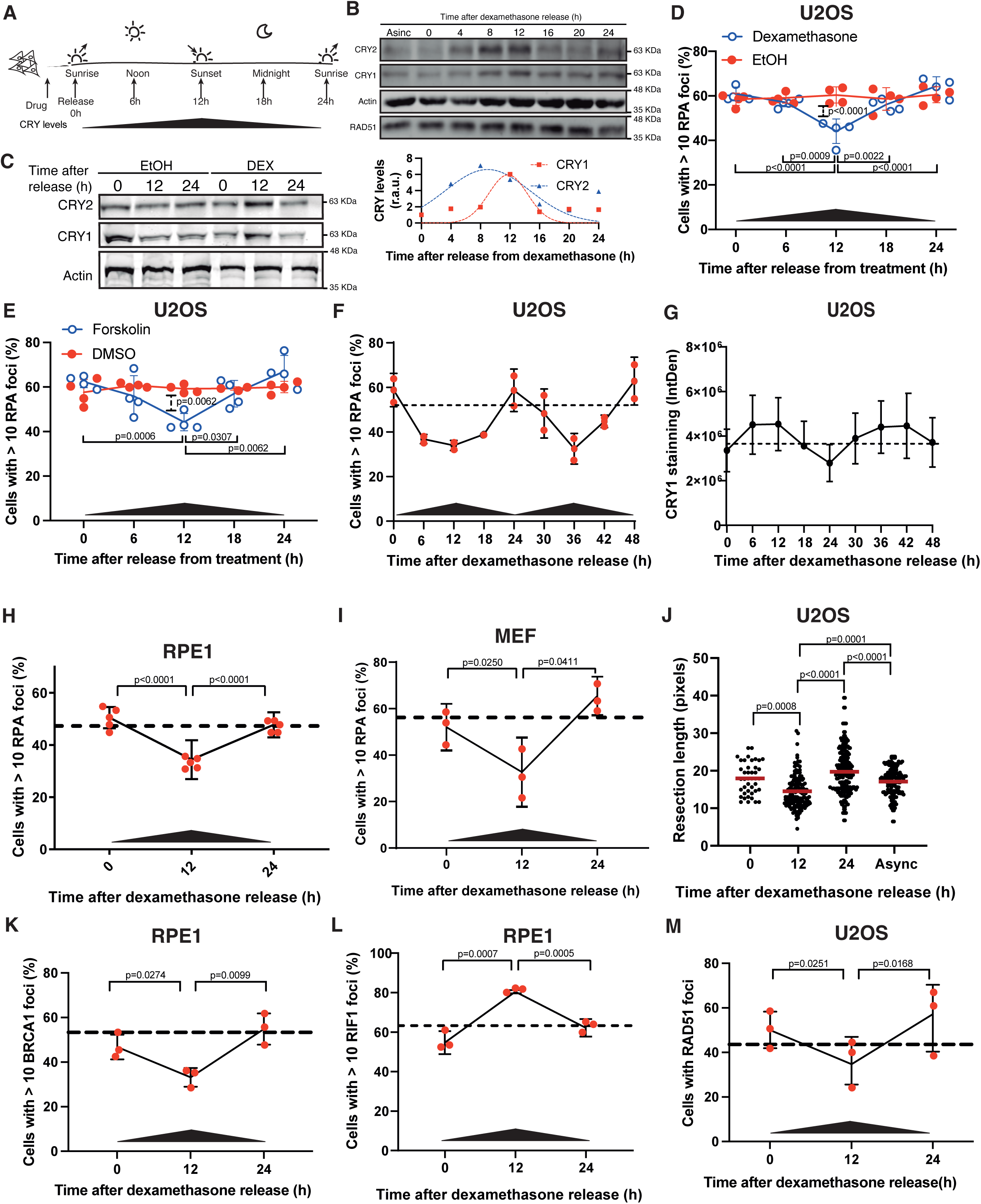
Resection oscillates following a circadian pattern. **A**, Schematic representation of the experimental setup for circadian synchronization. Cells were exposed to dexamethasone or forskolin for two hours. Then, cells were released by changing the medium, mimicking conditions with low CRY levels (i.e. early morning in human cells). Cells were then irradiated at the indicated timepoints that correspond to the depicted times of the circadian cycle, as expected by CRY levels of expression represented at the bottom by a dark triangle. **B**, U2OS cells exposed to dexamethasone for two hours were released by changing the medium. Protein samples were taken at the indicated timepoints, resolved in SDS-PAGE and blotted with the indicated antibodies. A representative image (top) and its quantification (bottom) are shown. CRY protein kinetics was inferred using a non linear fit gaussian model. **C**, Same as B but in cells released form DEX or EtOH as indicated. **D**, U2OS were synchronized as mentioned in A with either Dexamethasone (blue line) or EtOH as a control (red line) and irradiated with 10Gy at the indicated time points. 1 h later cells were immunostained for RPA. Then, the percentage of cells with more than 10 RPA foci was scored. At least 250 cells were analyzed for each sample in each replica. The average and standard deviation of four independent experiments is shown. Statistically significance was calculated using a two-way ANOVA. Only statistical different comparisons are depicted. CRY levels are represented with a black triangle at the bottom of the graph. **E**, Same as D, but using Forskolin (blue line) or DMSO as a control (red line). The average and standard deviation of four independent experiments is shown. **F**, Same as D, but cells were followed for 48 h after release from dexamethasone. **G**, Same as in F, but measuring CRY1 levels by immunofluorescence. **H**, Same as D but in RPE1 cells. Dashed line represents the value of an asynchronous control. The average and standard deviation of four independent experiments is shown. **I**, Same as H but in MEFs. **J**, The length of resected DNA was calculated as described in the methods section in cells synchronized and released from dexamethasone at the indicated times or in an asynchronous control. Each dot represents the length of an individual resected DNA fiber, and the median is shown in red. Statistical significance was calculated using a one-sided Mann–Whitney test. The experiment was repeated three times with similar results. **K**, Same as H but immunostaining for BRCA1. **L**, Same as K but immunostaining for RIF1. **M**, Same as K but immunostaining for RAD51 3 hours after DSB induction with IR in U2OS cells. Unless stated otherwise, for all main and supplementary figures, the average and standard deviation of three independent experiments are shown and p values are represented with one (p < 0.05), two (p < 0.01) or three (p < 0.001) asterisks. For the whole figure, source data are provided as a Source Data file.

Furthermore, this oscillation was also evident in the primary human cell line RPE1 (Fig. 1H, Suppl. Fig. 1F) and in mouse embryonic fibroblasts (MEFs; Fig. 1I, Suppl. Fig. 1G). Cell cycle is a major regulator of DNA end resection ^5,6,8,18^ and both the circadian and the cell cycles are known to crosstalk ^19^. However, our results could not be attributed to changes in cell cycle, as it was not affected by circadian synchronization (Suppl. Fig. 1H-J). To determine if the circadian rhythm controlled only the initiation of DNA end resection or if, instead, affected also the processivity of the process, we measured the actual length of resected DNA tracts at different positions on the circadian cycle (Fig. 1J). After release from synchronization, in agreement with a more permissive environment for resection, the length of DNA resected was longer than the asynchronous control. On the contrary, 12 hours, when CRY peaks, the extension of the resected DNA greatly diminished. Resection is controlled by multiple signals ^5,6^, chiefly the antagonistic relationship between the pro-resection BRCA1 and the anti-resection complex Shieldin ^20–22^. So, we then analysed the recruitment of BRCA1 and the Shieldin subunit RIF1 1h after the irradiation at different times of the circadian cycle. In agreement with our previous results, BRCA1 was more avidly recruited when DNA damage occurs right after release from synchronization whereas its localization was limited at the 12 hours timepoint, and then increase again 24 h after release (Fig. 1K; Suppl. Fig 1K). RIF1 showed the opposite recruitment pattern (Fig. 1L; Suppl. Fig 1L). Finally, breaks induced at the 12 hours timepoint were less likely to engage the recombinase RAD51 than those created at 0 or 24 h (Fig. 1M; Suppl. Fig 1M). Thus, we conclude that the circadian cycle in humans creates a more permissive environment for resection when CRY levels are low, but it is partially dampened when it peaks.

### DNA end resection reacts to CRY1 levels

We took advantage of our previously published results in which we tested the effect of depletion of all human genes in the balance between HDR and NHEJ ^9^. Interestingly, from the core and accessory circadian regulators, only CRY1 depletion skewed the balance between DSB repair pathways in a way that was compatible with our observations. Thus, we tested if resection was reacting directly to CRY1 levels. In agreement with a resection increase when the levels of this protein are naturally reduced, depletion of CRY1 stimulates the process and such effect was complemented by ectopic CRY1 expression (Fig. 2A; Suppl. Fig. 2A for depletion levels). Similar results were observed in RPE1 and MEFs (Suppl. Fig. 2B-C) and upon a CRISPR-mediated knockout of CRY1 in RPE1 cells (Suppl. Fig 2D). Strikingly, it is known that CRY1 loss does not abolish the circadian cycle, as its role is taken over by its paralog CRY2, but only shortens it by about 1 h ^23^. Thus, the fact that CRY1 depletion shows such a marked effect in resection stimulation strongly suggest that is specifically CRY1, and not the rest of the circadian built-in clock, what modulates this process. In agreement, CRY2 depletion did not cause an RPA foci defect whereas depletion of any PER paralogue showed a reduction in DNA end resection (Suppl. Fig. 2E), reinforcing this idea. But more importantly, CRY1 depletion not just delayed for 1h, but completely abolished the circadian oscillation of RPA foci formation (Fig. 2B; Supp. Fig. 2F). Interestingly, in the context of prostate cancer, it has been shown that CRY1 acts independently of the rest of the CRY-PER complex regulating DNA repair through controlling the stability of DNA repair factors ^24^. To confirm the effect of CRY1 depletion on RPA recruitment upon DSBs formation by different means, we took advantage of the DiVA system, in which AsiSI-mediated restriction creates breaks upon tamoxifen addition ^25^. ChIP-seq analysis of RPA recruitment on the best 214 AsiSI cutting sites, as previously described by BLISS ^26^, showed a stronger accumulation of RPA upon DSB formation in CRY1 depleted cells when compared with control cells (Fig. 2C, solid lines). As control, we compared what happens in 200 random sites in the genome, in which CRY1 depletion not only do not increase RPA presence but indeed reduces it (Fig. 2C, dashed lines).

**Figure 2.**
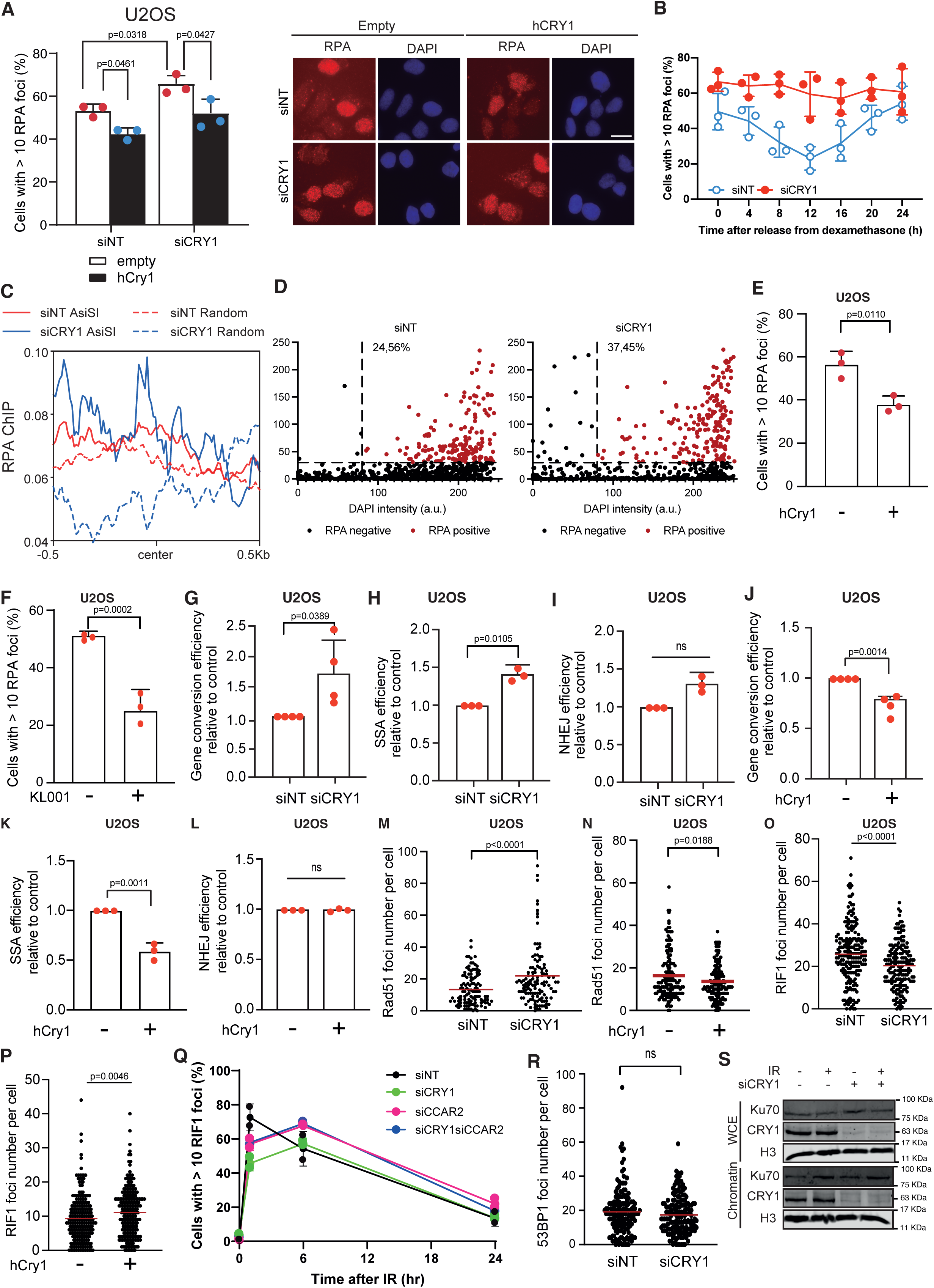
Resection oscillates following a circadian pattern. **A**, CRY1 (siCRY1) or control (siNT) depleted U2OS cells were transfected with the human version of CRY1 (black bars) or an empty vector (white). 1 h after irradiation with 10 Gy cells were immunostained for RPA. Then, the percentage of cells with more than 10 RPA foci was scored (left). The average and standard deviation of three independent experiments is shown. Statistically significance was calculated using a two-way ANOVA. Only statistical different comparisons are depicted. Representative images are shown on the right side. Scale bar 25 µm. **B**, Same as figure 1D, but in cells transfected with an siRNA against CRY1 or a control sequence, as indicated. Cells were irradiated every 4 hours after release in order to distinguish between a delay or a general upregulation of resection **C**, Average recruitment of RPA at the 214 best AsiSI cutting sites measured by ChIP-seq in samples transfected with an siRNA against CRY1 or a control sequence, as indicated (solid lines). One representative experiment, out of 2 with similar results, is shown. Similar analysis was performed in 200 random sites (dashed lines). **D**, QIBC representation of RPA foci formation. Cells treated as in A were immunostained for RPA. The intensity of RPA or DAPI were plotted in the Y and X axis, respectively. The vertical dashed line represents the threshold of DAPI intensity corresponding to S/G2 cells, whereas the horizontal line represents the threshold for RPA positive cells. The percentage of S/G2 cells showing RPA foci signal is indicated. A representative experiment out of 2 two independent replicates is shown. **E**, Same as A but in cells overexpressing or not CRY1 without siRNA transfection. The average and standard deviation of three independent experiments is shown. **F**, Same as A but in cells treated with the CRY1 stabilizer KL001. The average and standard deviation of three independent experiments is shown. **G**, Gene conversion was assessed in U2OS cells bearing the DR-GFP reporter in cells transfected with an siRNA against CRY1 (siCRY1) or a control sequence (siNT), as indicated. The amount of GFP positive cells, i.e., cells that have undergone repair by gene conversion, after induction with the nuclease I-SceI, were normalized to the siNT control taken as 1. The statistical significance was calculated using a Student’s t-test. The average and standard deviation of four independent experiments is shown. **H**, Same as G, but using the SSA reporter SA-GFP. **I**, Same as G, but using the NHEJ reporter EJ5-GFP. **J**, Same as G, but in cells bearing an empty vector or a vector harboring an ectopic version of CRY1. The average and standard deviation of four independent experiments is shown. **K**, Same as J, but using the SSA reporter SA-GFP. **L**, Same as J but using the NHEJ reporter EJ5-GFP. **M**, RAD51 foci 3 h after exposure to irradiation were scored by immunofluorescence in cells depleted for CRY1 (siCRY1) or control cells (siNT). The number of RAD51 foci per cell was counted and represented as a single dot. The median number of Rad51 foci is shown as a red line. Statistical significance was calculated using a Mann–Whitney test. A representative experiment out of three with similar results is shown. **N**, Same as M, but in cells transfected with a plasmid bearing an ectopic copy of CRY1 or an empty vector. **O**, Same as M, but immunostaining for RIF1. **P**, Same as N, but immunostaining for RIF1. **Q**, U2OS cells transfected with an siRNA against CRY1 (siCRY1), CCAR2 (siCCAR2), both (siCRY1siCCAR2) or a control sequence (siNT), were irradiated with 10Gy. Then, the percentage of cells with more than 10 RIF1 foci was scored at the indicated time points after irradiation. The average and standard deviation of three independent experiments is shown. **R**, Same as M, but immunostaining for 53BP1. **S**, Protein samples, depleted or not for CRY1 as indicated, were taken 1 h after irradiation. Upon cellular fractionation, both the whole cell extract (WCE) and the chromatin-bound fraction (Chromatin), were resolved in SDS-PAGE and blotted with antibodies against the indicated proteins. A representative experiment out of three replicas with similar results is shown. For the whole figure, source data are provided as a Source Data file.

Of note, this effect was underestimated, as CRY1 depletion increased the number of G1 cells (Suppl. Fig. 2G-H), a position on the cell cycle that is refractory to resection. To minimize this cell cycle effect, we repeated the immunofluorescence and analysed RPA foci determining the cell cycle phases by QIBC, using DAPI (DNA content) and EdU (DNA replication) as surrogates for cell cycle progression. CRY1 depletion increased the average number of RPA foci (above the horizontal dashed line) in S/G2 cells (right of the vertical dashed line) (Fig. 2D). Overexpression of CRY1 in cells transfected with a control siRNA reduced the amount of RPA foci upon DNA damage (Fig. 2A). We repeated the experiment in cells overexpressing human CRY1 but not transfected with any siRNA (Fig. 2E; Suppl. Fig 2I for representative images and 2J for expression levels) or treated with KL001, a drug that stabilizes CRY1 ^27^ (Fig. 2F; Suppl. Fig. 2K for expression levels and 2L for representative images). In both cases, resection was dampened. We conclude that we can artificially mimic the effect of the natural circadian oscillation in resection by either depleting CRY1 or overexpressing/stabilizing CRY1. We took advantage of this setup to test if DSB repair pathways efficiency were affected by CRY1 levels. Indeed, low levels of CRY1 stimulate both RAD51-dependent and -independent homologous recombination (Fig. 2G-H). Strikingly, this did not cause a reduction in NHEJ (Fig. 2I). Conversely, high levels of CRY1 hampered homology directed repair (Fig. 2J-K), but again did not affect the bulk of NHEJ (Fig. 2L). In agreement with this effect in recombination, RAD51 recruitment to DSB readily react to CRY1 levels (Fig. 2M-N and Suppl. Fig. 2M-N). The foci formation of the Shieldin complex 1 h after DNA damage induction was also affected by CRY1 expressions levels (Fig. 2O-P and Suppl. Fig. 2O-P), with an opposite pattern of RPA foci. In order to understand why the decrease in RIF1 formation might not be reflected in a decrease in NHEJ using the reporter (Fig. 2I) we performed kinetic study of RIF1 foci formation. We discovered that RIF1 recruitment was not really blocked, but delayed, in cells downregulated for CRY1 in a similar fashion to the depletion of the anti-resection factor CCAR2 (Fig. 2Q). This suggested that stimulating resection efficiency mostly delay NHEJ, but such changes do not reflect in a large reduction of the bulk NHEJ. In agreement, depletion of CRY1 does not abolish the recruitment of 53BP1 (Fig. 2R) or the bonafide NHEJ factor KU70 to damaged chromatin (Fig. 2S). Hence, it seems that only the anti-resection factor RIF1 is affected slightly by CRY1 levels, whereas the core NHEJ is unaffected. In order to understand if this nuances in RIF1 recruitment might contribute to the observed anti-resection phenotype, we decided to deplete RIF1 in the presence or absence of CRY1. Interestingly, even though CRY1 depletion slows down RIF1 accumulation (Fig. 2Q), this cannot explain the observed RPA defect as RIF1 depletion itself has very little effect (Suppl. Fig. 2Q). Thus, we conclude that despite this unsolved relationship between CRY1 and RIF1 foci formation, this is not relevant for the circadian-mediated regulation of DNA end resection and homologous recombination.

### CRY1 regulates the antagonist relationship between CtIP and CCAR2

We tested the effect of the depletion of the core resection factor CtIP in cells upon circadian synchronization. As expected, CtIP depletion reduced resection efficiency when compared with siRNA control transfected cells (Fig. 3A-B; Suppl. Fig 3A-B for statistics and Suppl. Fig. 3C for representative images). Strikingly, CtIP downregulation not only generally decreased resection, but mostly abolished the circadian oscillations of the process, suggesting that such regulation might rely on controlling this protein. In fact, 12 h after DEX release resection was limited to a degree compatible with an inhibition of CtIP.

**Figure 3.**
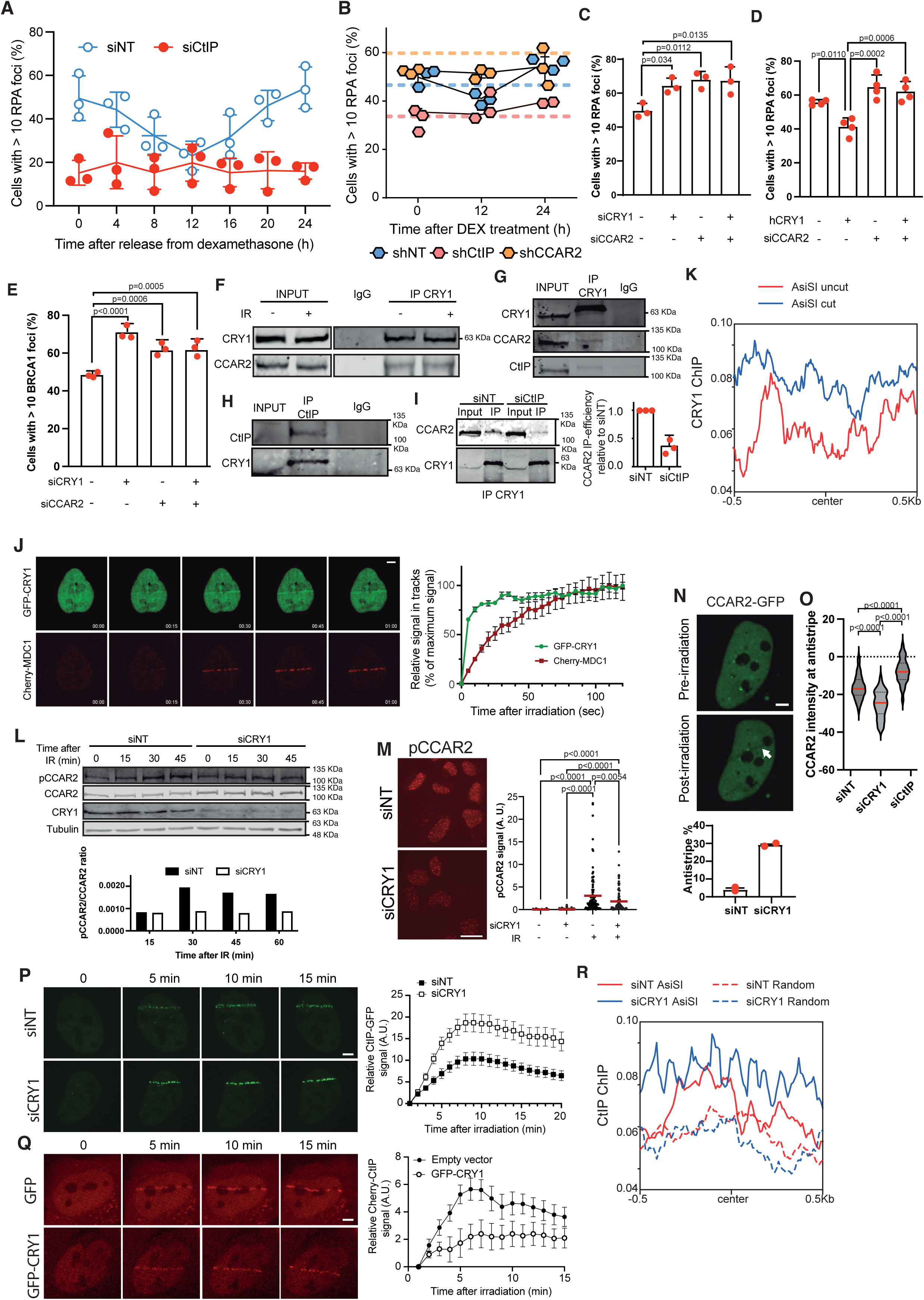
CRY1 levels control DNA end resection through CCAR2 retention at DSBs. **A**, U2OS transfected with siRNA against CtIP (siCtIP) or a control sequence (siNT), were synchronized and released as indicated in Figure 1A with dexamethasone. Cells were irradiated at the indicated time points and RPA foci were scored by immunostaining 1 h after irradiation. The average and standard deviation of three independent experiments is shown. Statistically significance was determined using a two-way ANOVA. **B**, Same as A but in cells bearing an shRNA against CCAR2 (shCCAR2), CtIP(shCtIP) or a control sequence (shNT). The average and standard deviation of three independent experiments is shown. **C**, RPA foci formation in U2OS cells transfected with the indicated siRNAs. A minus sign means transfection with a control sequence. The average and standard deviation of three independent experiments is shown. Statistically significance was determined using a one-way ANOVA. **D**, Same as C, but in cells transfected with a siRNA against CCAR2 or not and bearing a vector for CRY1 overexpression (hCRY1). Minus signs mean transfection with a siRNA control or the empty vector. The average and standard deviation of four independent experiments is shown. Statistically significance was determined using a one-way ANOVA. **E**, Same as C but immunostaining for BRCA1. The average and standard deviation of three independent experiments is shown. **F**, U2OS cells were irradiated with 10G (+) or mock treated (-). Protein samples were isolated in native condition and CRY1 was immunoprecipitated using a specific antibody against CRY1. IgG was used as immunoprecipitation control. Then, proteins were resolved in SDS PAGE and blotted with the indicated antibodies. A representative experiment out of three replicas with similar results is shown. **G**, Same as F but using an antibody against CRY1 for the IP. A representative experiment out of four replicas with similar results is shown. **H**, Same as G but using an antibody against CtIP for the IP. A representative experiment out of three replicas with similar results is shown. **I**, Same as F but in cells depleted for CtIP using an siRNA. The average and standard deviation of three independent experiments is shown. **J**, U2OS cells bearing Cherry-MDC1 and GFP-CRY1 constructs were laser microirradiated and imaged at the indicated times. Representative images of selected timepoints are shown on the right, and quantification of the signal, relative to the peak intensity taken as 100%, of 6 cells is represented on the right. Scale bar 5 µm. **K**, Average recruitment of CRY1 at the 214 best AsiSI cutting sites measured by ChIP-seq in samples exposed (cut, blue line) or not to (uncut, red line) to tamoxifen to induce a cleavage at AsiSI sites. **L**, U2OS cells transfected with an siRNA against CRY1 or a control sequence were irradiated with 10 Gy. Protein samples were taken at the indicated timepoints, and the chromatin fraction was purified. After resolving the sample in SDS PAGE, the amount of phosphorylated CCAR2 was determined using a specific antibody. Lamin A was used as a control. A representative western blot, out of three with similar results, is shown on top and its quantification of this specific western blot on bottom. **M**, U2OS cells treated as in L were immunostained using an antibody against phosphorylated CCAR2. Representative images are shown on the left side, and the quantification of the phospo-CCAR2 intensity per cell is plotted on the right side in cells irradiates (IR +) or not (IR −). A representative experiment out of three is shown. At least 50 cell per sample per experiment were analyzed. Statistical significance was analyzed using an unpaired one-sided Student’s T-test. Scale bar 25 µm. **N**, U2OS cells bearing a constitutively expressed CCAR2-GFP fusion were laser microirradiated. The percentage of cells showing a negative GFP staining 1 h after laser-microirradiated was quantified and plotted (right side) in cells depleted or not for CRY1. Representative image of a cell showing an anti-stripe (white arrow) before and 1 h after laser-microirradiation are also shown (left). The quantification of a representative experiment out of three is shown. At least 50 cell per sample per experiment were analyzed. Scale bar 5 µm. **O**, The intensity of CCAR2-GFP at the anti-stripe of cells showing them was calculated, setting the pan-nuclear intensity as 0, in U2OS cells laser-microirradiated upon depletion of CRY1 or CtIP. The average intensity in each case is shown as a red line. At least 70 cells with anti-stripes were analyzed in each sample. **P**, Recruitment of GFP-CtIP was measure upon laser microirradiation in U2OS cells transfected with an siRNA against CRY1 or a control sequence. Representative images are shown on the left side. Images were taken at the indicated times, and the intensity of GFP-CtIP at the laser-induced stripe was quantified and plotted. At least 15 cells were analyzed in each sample. Statistical significance was analyzed using an unpaired one-sided Student’s T-test. Scale bar 5 µm. **Q**, Same as P, but in cells stably expressing a Cherry-CtIP and transfected with a GFP-CRY1 plasmid or an empty vector. At least 15 cells were analyzed in each sample. Scale bar 5 µm. **R**, Same as figure 2C, but using and antibody against CtIP for ChIP. For the whole figure, source data are provided as a Source Data file.

CtIP is known to suffer a tight regulation at many different levels, including protein levels, mRNA stability, post-translational modifications, but also by the direct interaction with multiple proteins, including the antagonist factor CCAR2 ^7–9^. We reasoned that an inhibition of resection when CRY1 levels are high would agree with a peak of activity of this protein. Strikingly, CCAR2 is already linked with the regulation of circadian rhythms ^28,29^. CCAR2 depletion also abolished circadian oscillation of DNA end processing, but in this case allowing fully active resection at any given time of the circadian cycle (Fig. 3B; Suppl. Fig. 3A-B). Importantly, despite many circadian-mediated regulation happening by controlling the transcription of target genes, and a role of CRY1 regulating DDR factors expression in prostate cancer ^30^, neither CCAR2 nor CtIP protein levels change during the circadian cycle (Suppl. Fig. 3D). CCAR2 depletion was completely epistatic over both CRY1 depletion and CRY1 overexpression for RPA and BRCA1 foci formation, suggesting that CRY1 acts fully through stimulation of CCAR2 (Fig. 3C-E; Suppl. Fig. 3E-G). This epistatic effect cannot be attributed to large cell cycle changes (Suppl. Fig. 3H). This fits with the idea the circadian clock affects resection and recombination directly through a specific activity of CRY1, and not through the normal core clock function. Thus, we wondered if CRY1 protein might directly interact with CCAR2. Indeed, both proteins physically interact constitutively, i.e. regardless of the exposure to IR (Fig. 3F). CCAR2 and CtIP also interact constitutively ^9^. Indeed, we could co-immunoprecipitate CtIP and CRY1 by reciprocal IPs (Fig. 3G-H) and detect their proximity by Proximity Ligation Assay (PLA; Suppl. Fig. 3I). Furthermore, CtIP depletion reduced the interaction between the other two proteins (Fig. 3I), suggesting that all three proteins might interact together.

Considering this interaction, we wondered if CRY1 was recruited to sites of chromosome breaks as it has been described for both CtIP and CCAR2 ^9^. Using a GFP-CRY1 we observed a fast accumulation of this protein at DSBs by laser microirradiation with an even faster kinetic than the early DDR factor MDC1 (Fig. 3J; Suppl. movies 1 and 2). Strikingly, in agreement with a specificity of CRY1 but not CRY2 in regulating DNA end resection, only the former but not the later was recruited to AsiSI sites after a DSB was induced by cleavage with the enzyme (Fig. 3K; Suppl. Fig 3J).

We previously showed that CCAR2 interacts with CtIP, and upon the appearance of DSBs controls CtIP recruitment and spreading from the breaks ^9^. In less permissive conditions for resection, CCAR2 is maintained at DSBs, where is phosphorylated by ATM to block the DNA end processing ^9^. On the contrary, in resection-proficient conditions it is evicted from the DNA ^9^. Considering our data, we wondered if CRY1 could be regulating CCAR2 retention/eviction from sites of damaged chromatin. Strikingly, phosphorylated CCAR2 recruitment/retention was diminished if CRY1 was downregulated, as observed by the fraction bound to chromatin upon DNA damage induction (Fig. 3L) and retained by immunofluorescence upon pre-extraction (Fig. 3M). Then, we checked CCAR2 eviction from the vicinity of the damaged DNA by laser microirradiation, that can be observed as a negative signal upon known as an anti-stripe ^9^. Interestingly, CRY1 depletion increases the formation of CCAR2 anti-stripes 7-fold (Fig. 3N, see white arrow for an example of a laser-induced anti-stripe). Furthermore, not only the number of cells showing antistripes was increased upon CRY1 depletion, but quantification of the remaining signal at the laser-induced antistripes showed a stronger reduction of the CCAR2 signal (Fig. 3O), agreeing with a reduced capacity to retain the protein there. On the contrary, when CtIP was downregulated even those fewer cells that showed antistripes still maintain a significant proportion of the GFP-CCAR2 signal, arguing that there is some CCAR2 eviction, but a considerable fraction of the protein is still retained (Fig. 3O). Then, we tested how CtIP recruitment/retention to laser lines responded to CRY1 levels live. Cells transfected with a control siRNA accumulate quickly, within minutes, GFP-CtIP laser stripes (Fig. 3P, Suppl. movie 3). However, when CRY1 was depleted, we observed that GFP-CtIP accumulation was stronger and at a faster rate (Fig. 3P and Suppl. movie 4). The opposite effect was observed upon CRY1 overexpression using a GFP-tagged version of the protein, that causes a reduction on the recruitment of Cherry-CtIP at laser lines (Fig. 3Q; Suppl. movies 5 and 6). Furthermore, an increased retention of CtIP at sites of AsiSI-induced breaks could also be observed upon CRY1 depletion by ChIP-Seq at AsiSI cleavage sites but not at random genomic positions (Fig. 3R). Thus, CRY1 levels control the retention or eviction of CCAR2, a requisite to unleash CtIP and allow DNA end resection.

### Resection is modulated by DNA-PK-mediated phosphorylation of CRY1

Our data fit with a model in which CRY1 presence reinforces the antagonistic effect of CCAR2 over CtIP, stabilizing CCAR2 presence on damaged chromatin hence blocking CtIP release and, therefore, resection. However, and differently of the cell cycle regulation of resection, this is not an all or nothing mechanism. So, we hypothesized that even when CRY1 levels are high there must be a mechanism that allows some resection. Thus, we decided to analyse how the DDR finetunes the circadian oscillations of DNA end resection. Previously, CRY1 itself has been shown to be a substrate of the kinase DNA-PK ^31^, a modification conserved in mammals but no other vertebrates. Interestingly, these phosphorylation sites are absent in CRY2. In fact, CRY1 directly physically interact with both DNA-PKcs and KU70 according to mass spectrometry data ^32^.Considering its anti-resection and pro-NHEJ role, it made sense that such phosphorylation stimulates CRY1-mediated inhibition of CtIP-mediated resection. So, we first tested CRY1 phosphorylation upon treatment with IR in control cells or cells depleted (Suppl. Fig. 4A) or inhibited (Suppl. Fig. 4B) for DNA-PKcs. Indeed, we could see an IR-induced phosphorylation of CRY1, and such band was reduced by DNA-PK inhibition or depletion. However, some residual phosphorylation was still present on those conditions. Thus, it is clear that there are additional kinases involved in CRY1 phosphorylation upon DNA damage, but their nature and relevance should be addressed in follow up projects.

Thus, we analysed the effect of such modification on the modulation of DNA end resection and recombination. Indeed, ectopic expression of human CRY1 or its stabilization with KL001 hampered DNA end processing only if DNA-PK was active (Fig. 4A-B; Suppl. Fig. 4C-D for representative images). Furthermore, DNA-PK inhibition abolished the resection dampening observed 12h after release from DEX (Fig. 4C). CRY1 has been shown to be phosphorylated at three specific sites ^33^. So, to test if this effect was direct, we checked the effect of the overexpression of a CRY1 version that cannot be phosphorylated by DNA-PK (CRY1-3A) or that mimicked constitutive phosphorylation (CRY1-3E) in resection in the presence or absence of DNA-PK inhibitor. As shown in figure 4D, overexpression of wildtype CRY1 hampered resection only when DNA-PK was active, in agreement with a requirement of DNA-PK-mediated phosphorylation for its anti-resection role (see Suppl. Fig. 4E for representative images). In stark contrast, a CRY1 that cannot be modified by DNA-PK did not impair resection even when DNA-PK was functional. On the contrary, a constitutive phosphorylation of CRY1 was enough to hamper resection even in the presence of DNA-PK inhibitor. To determine the molecular role of those phosphorylations, we measured CRY1 accumulation at laser-lines in real time in the different mutant backgrounds (Figure 4E; Suppl. movies 7-9). All three versions of the protein were recruited to DSBs with similar kinetics. As a control, we confirmed that all three constructs are expressed to similar levels and maintain a comparable protein stability to the endogenous CRY1 (Suppl. Fig. 4F).

**Figure 4.**
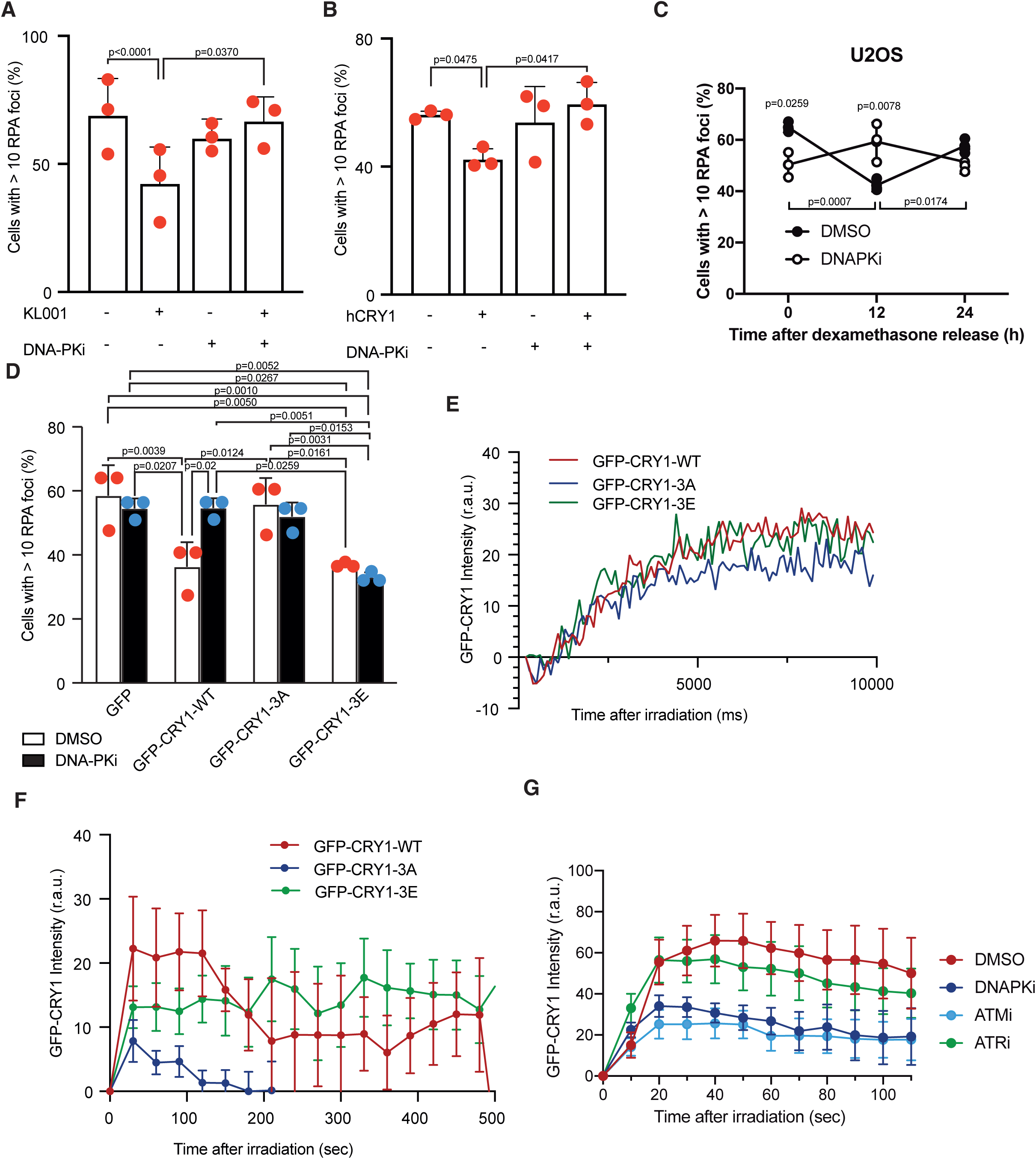
DNA-PK-dependent phosphorylation of CRY1 modulates resection. **A**, U2OS cells were exposed to the CRY1 stabilizer KL001 and/or the DNA-PK inhibitor NU7441 as indicated with the plus and minus signs, and then irradiated and immunostained for RPA. The average and standard deviation of three independent experiments is shown. Statistically significance was determined using a one-way ANOVA. **B**, Same as A but in cells transfected with an ectopic version of CRY1 or the empty vector instead of exposed to KL001. The average and standard deviation of three independent experiments is shown. Statistically significance was determined using a one-way ANOVA. **C**, U2OS cells treated with an inhibitor of DNA-PK or DMSO as a control were treated as in Figure 1B. Other details as Figure 1B. The average and standard deviation of three independent experiments is shown. Statistically significance was determined using a one-way ANOVA. **D**, Cells stable transfected with GFP-fusions of wildtype CRY1 (GFPCRY1-wt), a non-phosphorytable version of CRY1 (GFPCRY1-3A), a phospho-mimicking version (GFPCRY1-3E) or the empty vector (GFP) were treated with NU7441 or not, as indicated, irradiated and immunostained for RPA. The average and standard deviation of three independent experiments is shown. Statistically significance was determined using a two-way ANOVA. **E**, U2OS stably expressing GFP-CRY1 versions were laser microirradiated and imaged at the indicated times. For each timepoint, the average intensity of at least five cell images is plotted. **F**, Same as E, but for longer time points. **G**, Same as in F, but with cells treated with inhibitors for DNA-PK, ATM, ATR or DMSO as a control, as indicated.

Then, we tested if retention of CRY1 was affected by DNA-PK-mediated phosphorylation by analysing laser lines with longer time courses (Figure 4F; Suppl. movies 10-12). Strikingly, we observed that albeit all three CRY1 variants were readily recruited, only when CRY1 was phosphorylated it was retained at sites of damaged DNA. Thus, these data suggest that upon the appearance of DSBs CRY1 is recruited to sites of broken chromosomes, but only when it is phosphorylated by DNA-PK it is retained. We wondered if other PIKKs could affect CRY1 accumulation directly or indirectly. Indeed, as expected, DNA-PK inhibition affects CRY1 retention similarly to the CRY1-3A mutant (Fig. 4G). Similar effects were obtained upon depletion of DNA-PKcs with an siRNA, discarding that the inhibitor effect was due to the enzyme trapping (Suppl. Fig. 4G). Additionally, the inhibition of ATM, but not ATR, also diminished CRY1 retention (Fig. 4G). If this reflects the effect of ATM on CCAR2 retention or others, more indirect, effects, remains to be studied. Of note, neither DNA-PKcs nor ATM nor ATR levels change during a circadian cycle nor in response to CRY1 depletion (Suppl. Fig. 4H).

### Circadian regulation of DSB repair affects cell survival to genotoxic agents

Because of CRY1 regulating DNA end resection and, therefore, HDR, cells become hyper-resistant to several DNA damaging agents when this factor was depleted (Fig. 5A-B) and hyper-sensitive when it was stabilized using KL001 (Fig. 5C-D). Albeit this effect might seem modest, it is of a similar level to that observed upon depletion of core resection factors such as CtIP ^34^. In order to determine if this was only observed upon natural oscillations of CRY1, we repeated the clonogenic assays in cells synchronized with DEX and releases for different times. Indeed, as expected, whereas at 0 and 24 hours after release the survival was high, at 12 hours, coinciding with a peak in CRY1 and a reduction in RPA and RAD51 foci, survival was reduced (Fig. 5E). Thus, survival to genotoxic agents was also affected by natural, circadian, oscillations of CRY1 levels. So, we conclude that HDR stimulation when CRY1 was low facilitated repair, whereas high CRY1 induced a reduction in HDR that hampered DNA repair. Indeed, CRY1 depletion reduced the number of cells with micronuclei 24 h after irradiation with 10 Gy (Fig. 5F; see Suppl. Fig. 5A for representative images) and the number of unrepaired breaks in the next mitosis upon DSB induction (Fig. 5G left; see Suppl. Fig. 5B for representative images). On the contrary, stabilization of CRY1 increases the number of unrepaired breaks in the following mitosis after exposure to IR (Fig. 5G right; see Suppl. Fig. 5C for representative images). Moreover, when we measured the disappearance of γH2AX 24h after irradiation as a proxy of DSB repair completion, we observed that cells depleted of CRY1 were more efficient than control cells (Fig. 5H; see Suppl. Fig. 5D for representative images). To confirm that CRY1 acts through CCAR2, we tested survival to IR and Etoposide upon single and double depletions (Fig. 5I-J). As expected, both single depletions affected survival similarly, and double depletion showed an epistatic effect. The same genetic relationship was seen on micronuclei formation (Fig. 5K). Furthermore, we have directly tested for the persistence of DSBs by COMET assay. Indeed, depletion of either CRY1 or CCAR2 strongly reduced the tail length, indicating a faster elimination of the breaks (Fig. 5L, Suppl. Fig. 5E). As expected for factors involved in the same pathway, double depletion of both did not render any further effect. Thus, we conclude that the levels of CRY1 are important to maintain the stability of the genome through its regulation of CCAR2.

**Figure 5.**
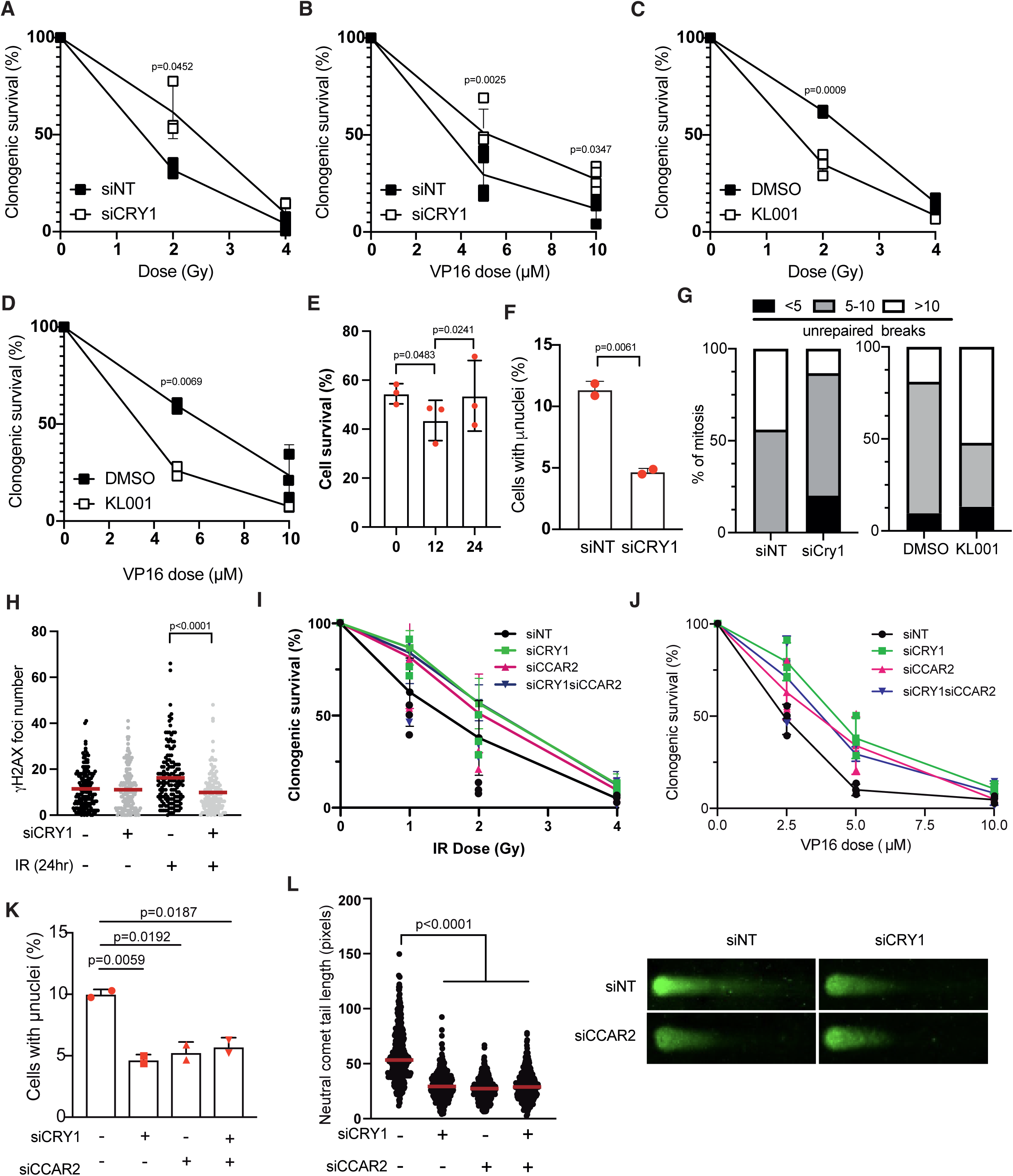
CRY1 levels modulate the response to DNA damaging agents *in vitro*. **A**, U2OS transfected with an siRNA against CRY1 or a control sequence were seeded at low density and irradiated at the indicated dose. Cells were left to grow for 10 days and the survival fraction, compared with an unirradiated sample taken at 100%, is plotted. The average and standard deviation of three independent experiments is shown. Statistically significance was determined using a one-way ANOVA. **B**, Same as A but in cells treated with the indicated doses of etoposide (VP16) for 1 hour. The average and standard deviation of three independent experiments is shown. Statistically significance was determined using a one-way ANOVA. **C**, Same as A but in cells pre-exposed to the CRY1 stabilizer KL001 or DMSO. The average and standard deviation of three independent experiments is shown. Statistically significance was determined using a one-way ANOVA. **D**, Same as B but in cells pre-exposed to the CRY1 stabilizer KL001 or DMSO. The average and standard deviation of three independent experiments is shown. Statistically significance was determined using a one-way ANOVA. **E**, Same as A, but in cells synchronized with DEX, released for the indicated time and exposed to 2 Gy of IR. The average and standard deviation of three independent experiments is shown. Statistically significance was determined using a one-way ANOVA. **F**, U2OS cells depleted or not of CRY1, as indicated, were irradiated with 10 Gy. 24 h after irradiation cells were stained with DAPI and the percentage of cells with micronuclei was calculated. The average and standard deviation of two independent experiments is shown. Statistically significance was determined using a one-way ANOVA. **G**, Percentage of chromosome breaks present in mitotic cells upon irradiation with 2 Gy transfected with the indicated siRNAs (left) or exposed to KL001 or DMSO (right). **H**, Number of γH2AX foci in U2OS cells unchallenged (-) or 24 h after irradiation (+; 10 Gy), in cells transfected with an siRNA against CRY1 (+) or a control sequence (-), as indicated. A representative experiment out of three is shown. Statistical significance was analyzed using an unpaired one-sided Student’s T-test **I**, Same as A but in cells depleted for CRY1, CCAR2 or both proteins simultaneously. The average and standard deviation of three independent experiments is shown. **J**, Same as B but in cells depleted for CRY1, CCAR2 or both proteins simultaneously. The average and standard deviation of three independent experiments is shown. **K**, Same as F but in cells depleted for CRY1, CCAR2 or both proteins simultaneously. The average and standard deviation of three independent experiments is shown. Statistically significance was determined using a one-way ANOVA. **L**, Cells treated as in panel K were assayed for DNA break presence using neutral comet assay. Left, quantification of one out of three similar experiment. Each dot represents one nucleus. Right, a representative image of a nucleus for each condition is shown. A representative experiment out of three is shown. Statistical significance was analyzed using an unpaired one-sided Student’s T-test. For the whole figure, source data are provided as a Source Data file.

### CRY1 levels affects tumour biology and prognosis

Circadian proteins are commonly deregulated in tumour samples. Thus, we wondered if such relationship with the repair of induced DSBs and genomic instability was maintained in those pathological setups and, furthermore, if it could be exploited in the clinic. First, we reasoned that if CRY1 levels impact genomic instability it might affect the probability to acquire secondary mutations in cancer samples that lead to a new tumour event. Using Pan-Cancer data from The Cancer Genome Atlas (TCGA) we observed that cells with high levels of CRY1, i.e., with a decreased recombination capacity and worse repair according to our model, showed an acceleration in the appearance of a new tumour event (Fig. 6A). According to our data, higher levels of CRY1 were associated with an increased exposure to the mutational signature associated with defective HDR (SBS3; Fig. 6B), i.e. the one defined by BRCA1 or BRCA2 absence. In fact, tumours with low levels of CRY1 present virtually no exposure to this specific mutational signature. Then, we reasoned that despite this increased mutagenicity of cancer samples with high levels of CRY1, this reduced repair ability can be exploited by using therapeutic agents designed to create DSBs. Indeed, using again TCGA data, we observed that breast cancer patients with tumours expressing higher CRY1 levels were more responsive to radiotherapy, in agreement with the idea that high CRY1 levels render the tumour cells more sensitive to radiation (Fig. 6C). On the contrary, breast tumours with lower CRY1 expression were less efficiently treated with radiotherapy (Fig. 6C), with a decrease in the median survival of 1.5 years (571 days). Strikingly, similar results were observed regarding CCAR2 levels (Fig. 6D), albeit with an even bigger increase in median survival of over 2.5 years (957 days), reinforcing that both CCAR2 and CRY1 levels affect similarly cancer prognosis. Furthermore, in heterotopic, subcutaneous, xenografts models using HCT116 cells bearing a constitutive shRNA against CRY1 or a control sequence, we observed a faster growth of tumours lacking CRY1 and a better tolerance to etoposide treatment (Fig. 6E, Suppl. Fig 6). Indeed, in samples expressing CRY1, etoposide treatment caused a 25% reduction in tumour growth rate. In agreement with a mild increase in HDR, and a reduced cytotoxic effect of etoposide, such decrease in tumour growth rate was slightly reduced to 18% in samples lacking CRY1. This may suggest that CRY1 loss is associated with tumour evolution and sensitivity to treatment *in vivo*.

**Figure 6.**
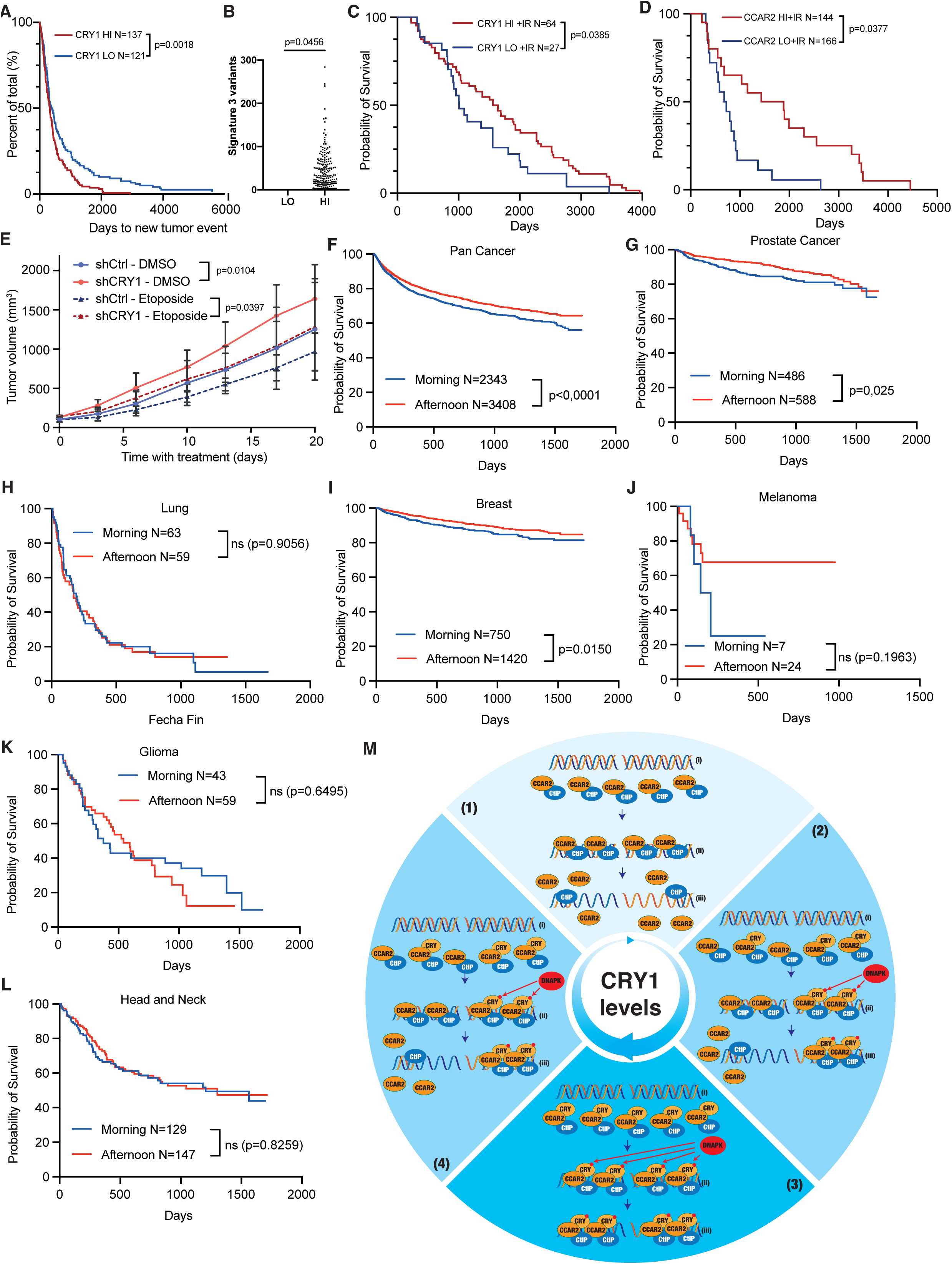
CRY1 levels modulate the behaviour of cancer cells and their response to treatment. **A**, Elapsed days for a second tumour event in cells stratified for its CRY1 levels in two categories: High (red line) or low (blue line). Data from TCGA. The number of tumour samples is indicated. Statistical significance was analyzed using Kaplan-Meier test. **B**, Pan-cancer data from the TCGA were separated according the levels of CRY1 and the appearance of mutations associated with the mutational signature 3, caused by defective HDR, was scored and represented. Statistical significance was analyzed using an unpaired one-sided Student’s T-test**. C**, Survival of breast cancer patients treated with radiotherapy according the CRY1 levels of the tumour cells. Data from TCGA. Other details as in A. Statistical significance was analyzed using Kaplan-Meier test. **D**, Same as C but regarding CCAR2 levels. Statistical significance was analyzed using Kaplan-Meier test. **E**, Nude mice were grafted with HCT116 cells depleted for CRY1 on the right flank and control cells on the left flank. Mice were treated with etoposide as indicated in the method section. The average and standard deviation of tumour volume of 6 mice for each condition and timepoint are plotted. Statistical significance of the slopes after linear regression of the data is shown. **F**, Overall survival of cancer patients treated primarily with radiotherapy at the Radiotherapy Service of the University Hospital Virgen Macarena considering if they were irradiated in the morning (blue line) or the afternoon (red line). Statistical significance was analyzed using Kaplan-Meier test. **G**, Same as F but considering only prostate cancer patients. Statistical significance was analyzed using Kaplan-Meier test. **H**, Same as F but considering only lung cancer patients. Statistical significance was analyzed using Kaplan-Meier test. **I**, Same as F but considering only breast cancer patients. Statistical significance was analyzed using Kaplan-Meier test. **J**, Same as F but considering only melanoma patients. Statistical significance was analyzed using Kaplan-Meier test. **K**, Same as F but considering only glioma patients. Statistical significance was analyzed using Kaplan-Meier test. **L**, Same as F but considering only head and neck cancer patients. Statistical significance was analyzed using Kaplan-Meier test. **M**, Schematic representation of CRY1-mediated regulation of DNA end resection regarding CRY1 levels, represented for the circular arrow at the center of the image. (1) When CRY1 is low, CtIP and CCAR2 are recruited to DSBs (ii). However, CCAR2 cannot be maintained into chromatin, releasing CtIP and promoting maximum resection (iii). (3) On the contrary, when CRY1 peaks, it forms a complex with CCAR2. Upon DSB formation (i), all three proteins are recruited to damaged chromatin, where CRY1 is phosphorylated by DNA-PK (ii). This locks CRY1 and, consequently, CCAR2 on chromatin, hampering CtIP activity (iii). During the rest of the cycle (2 and 4), when CRY1 levels fluctuate, an intermediate situation ensues. I. e. some breaks will be promptly resected ((iii) left) and some will not ((iii) right), depending on the levels of CRY1.

Then, we reasoned that if CRY1 levels affected the response to genotoxic agents such as radiotherapy, it is possible that the time of the day when patients are irradiated might impact the efficiency of this treatment. In a retrospective analysis using patient data from the Radiotherapy Service from the University Hospital Virgen Macarena (Seville, Spain) we analysed this response stratifying the patients in two groups: those irradiated in the morning versus those irradiated in the afternoon/evening. Using data from all cancer types (pancancer) we observed a significant difference between those two groups (Fig. 6F). In agreement with our CRY1 levels results, irradiation in the afternoon-evening, when CRY1 levels build up, rendered tumour samples more sensitive to radiotherapy and improved patient prognosis. We hypothesize that this effect might be small because many cancers show deregulation of circadian proteins, and in this case, patients might not always benefit from this association between therapy and time of the day, the so-called chronotherapy^35^. For example, directly deregulation of CRY1 has been observed in some tumours such as lung cancer, whereas in other tumour types like prostate cancer deregulation of circadian rhythms does not rely in changes in CRY1 levels ^36^. Strikingly, careful examination of the data agreed with this idea and, indeed, whereas prostate cancer patients do benefit of irradiation at specific times of the day (Fig. 6G), lung cancer patients did not (Fig. 6H). This differential behaviour can be observed also in other cancer types. For example, and in agreement with our TCGA analysis for CRY1 levels (Fig. 6B), a benefit in an afternoon irradiation can be seen in breast cancer patients (Fig. 6I). An even stronger effect is observed in melanoma, maybe reflecting a response to the direct exposure to sunlight of the tissue, but the low number of patients with this type of cancer that are treated with radiotherapy precludes the statistical significance of the differences (Fig. 6J). On the contrary, other tumour types such as head and neck or gliomas do not benefit from a chronotherapeutic approach (Fig 6K-L).

## DISCUSSION

Circadian rhythms and the repair of DNA damage are strongly evolutionary linked. Indeed, CRY1 belongs to the Cryptochrome family, in which bacterial photolyases are included ^37^. Similarly, Timeless acts as a bona fide circadian accessory factor in Drosophila, whereas in mammals has shifted its role toward a checkpoint function ^38^. Strikingly, in Drosophila there is a second family member, Timeout, that has been related to the repair of DSBs ^38^. So, it seems that during evolution a bleeding between circadian-repair proteins has occurred multiple times. Although this simple crossing between circadian-repair pathways might explain why in mammals CRY1 behaves as both a circadian and a repair factor, a more intriguing hypothesis is that this dual nature reflects an evolutionary benefit of synchronizing DSB repair with the night/day cycle. So, a likely explanation is that this regulation prepares cells to face a load of DSBs at specific times of the day. Considering that UV light is a strong inductor of DNA damage, including DSBs when not repaired by NER, this connection might not seem surprising. Indeed, NER has been clearly shown to be regulated by the circadian clock, attuning this repair pathway to the part of the day in which UV-derived DNA lesions will appear ^39^. So, a possibility is that this connection reflects a need to prepare cells to face unrepaired UV-lesions that can lead to DSBs that are more likely to appear during the day. However, we do not favour this idea, mostly by two reasons. First, most mammalian cells do not suffer this kind of damage as they are protected from the mutagenic effect of UV by the skin. And second and more importantly, we have observed a conservation of the role of CRY1 in mice that disagrees with this hypothesis. Relevantly, mice are nocturnal animals, so they have a 12-hour shift in their circadian clock respect to humans and other diurnal mammals, with CRY1 peaking at early morning and sporting a maximal reduction at evening ^40^. Thus, in mice cells recombination peaks at the equivalent of dusk in an ideal cycle and it is reduced at dawn, opposite to what happens in human cells. So, for mice to maintain the same CRY1-dependent circadian regulation points out toward a different source of DNA damage that does not appear during the day but accumulates at night. An alternative explanation will be replication-born damage. It has been established that the circadian clock and the cell cycles are in phase, and in human cells that means that replication takes usually place at specific times of the day ^41^. Albeit it is not completely clear how this coordination works, and the time of the day in which replication takes place varies among different cell types, it has been proposed that replication only takes place at specific times (gates) of the day in specific circadian windows ^41^. So, it is possible that the regulation of resection by CRY1 levels is a way to reinforce the cell cycle regulation of homologous recombination, ensuring that resection and HDR only happen in specific circadian windows too. A better understanding on the synchronization between replication and the circadian clock is required to test this idea. However, considering all our data together, we favour a different explanation. Indeed, we propose that this circadian synchronization might prepare mammalian cells to a source of damage that is more likely to appear during the day in humans and at night in mice. So, we hypothesize that this circadian regulation might reflect a need to attune DSB repair with the appearance of damage caused by the cellular metabolism, another well-stablished source of DNA damage ^42^. Interestingly, CCAR2 has been proposed to coordinate the cell metabolism with the circadian clock through its interaction with the Rev-erbα receptor ^43^, suggesting that it might mediate also further coordination of these two aspects with DSB repair. This will then be more related to CRY1 peaks and valleys that the actual night or day. In this model, during the more active phase of the cells, i.e., when the metabolism is more active, higher rates of recombination will be required. So, in humans, cells will benefit of a boost of recombination at early morning whereas mice will favour such increase at evening as a preparation of the initiation of the active phase of their circadian cycle. Then, during this more active phase recombination can slowly decline until dusk (in human) or dawn (mice) in readiness for a more reduced metabolic status, meaning that the chances of encounter DSBs that require recombination for repair are reduced. Then, the cells will increase slowly again their ability to activate resection to face the next day/night cycle.

But how does this circadian regulation impinge on the regulation of DNA end resection? We favour a model in which the regulation does not rely on the whole circadian clock, but rather it is an indirect effect through the circadian oscillation of a specific factor, CRY1, that independently of its other circadian roles directly controls resection through CCAR2 and its inhibition of CtIP. Strikingly, this cryptochrome paralogue specificity might reflect the lack of conservation of the C-terminal tail between CRY1 and CRY2, including the absence of the DNA-PK phosphorylation sites in the later. This align with the idea that such phosphorylation is required to lock CRY1 at sites of DSBs, explaining the lack of recruitment of CRY2 to such places. In any case, we do not discard other alternative mechanisms in other animals that lack this specific phosphorylation or even CRY1 itself. In our model (Fig. 6M) in an ideal circadian cycle, in which low and high levels of CRY1 alternates every 12 h, resection will follow such oscillations. In this scenario, we envision that the levels of CRY1 are critical to retain CCAR2 and restrain CtIP by a physical interaction between these three proteins. Indeed, we think that at when CRY1 is low (Figure 6M, 1) CCAR2 and CtIP are recruited together to DSBs as previously proposed ^44^ (Fig. 6M, 1, (ii)). But in the absence of CRY1, CCAR2 cannot stay on damaged DNA, so it is readily evicted from chromatin unleashing CtIP and creating a pro-resection environment (Fig. 6M, 1, (iii)). On the contrary, when CRY1 peaks (Figure 6M, 3), CRY1 will be constitutively interacting with CCAR2. Therefore, when a DSB appear all three proteins will be recruited on chromatin in a very rapid response (Fig. 6M, 3, (ii)). Then, the activation of DNA-PK will cause the phosphorylation of CRY1, what will lock it on DNA, fastening CCAR2 also on the broken chromatin and constraining CtIP activity (Fig. 6M, 3, (iii)). This is also reinforced by ATM activity, either by affecting CtIP or CCAR2 or by additional mechanisms. During either the increase or decline of CRY1 levels (Figure 6M, 2 and 4), cells will react differently depending on the actual amount of this circadian factor. Interestingly, this first regulation represents only a first layer of the modulation of recombination by the circadian proteins. Interestingly, Legube and colleagues have described that PER proteins affect homology mediated repair through a non-canonical function independent of others circadian core clock components ^45^. Specifically, they see that PER2 reduces the clustering of DNA double strand break by helping their localization at the nuclear envelope, therefore fostering the strand invasion step of recombination and minimizing genetic translocations. Due to an uncoupling on the oscillation of CRY1 and PER proteins, this mechanism could reinforce a reduction of recombination during the night, collectively supporting a two-tier model in which resection is dampened in humans following CRY1 peak, and then even those breaks that have been partially resected will later fail to engage a homologue sequence due to reduce PER levels later on. Additionally, it has been recently proposed that BMAL1-CLOCK stimulate resection and recombination through the acetylation of histone H4 ^46^, hence suggesting a further stimulation of recombination during the day. Importantly, this regulatory mechanism will impact in the quality of repair, with a strong effect in the appearance of genomic translocation when defective. Of note, a previous report discovered that in the context of prostate cancer, the androgen receptor stimulates CRY1, that then affects DNA repair by regulating the expression of DDR ^24^. However, they find the opposite role of CRY1 and propose that stimulates recombination. If this is due to differences in cell types or a consequence of hormone induction remains to be resolved.

This built-in mechanism(s) will act at the cellular level. Additionally, there is a secondary control of the circadian rhythms at the level of light sensing that can reset the cellular clock, that in the case of the mammals require the detection of light by the eyes, the processing of this signal at the suprachiasmatic nucleus (SCN), and then the transmission of this information to all the cells in the organism ^47^. How this supracellular regulation impinges in this regulatory network we have uncovered should be explored in the future. Indeed, our data suggest that perturbations of the circadian rhythms might have consequences in the ability of the cells to repair broken chromosomes. Furthermore, it will affect not only their ability to repair DSBs, but also the balance between the more error-free HDR and the error-prone NHEJ, hence putatively impacting both the quantity and quality of the repair. This might, in turn, affect the stability of the genome, altering the mutational load. Indeed, we see an increased exposure to the mutational signature associated with defective HDR in cells with high levels of CRY1. Strikingly, it has been known for decades that people that alternate between different day-night shifts at work or suffering of chronic jetlag are more prone to develop cancer and they generally have a poorer prognosis ^48,49^, albeit the reasons are far from clear. One tantalizing idea is that these workers have a deregulation between their DSBs repair pathways due to the circadian perturbations, so they have a higher chance toward a mutagenic repair that might contribute, among other factors, to tumour development. Along the same lines, the same explanation can be invoked to understand why circadian regulators are often altered in many cancer cells ^50,51^. So, tampering with the built-in circadian clock, either by external effects, such as shift changes, or internal causes, such as mutation of circadian genes, will contribute to an increase in the mutagenic burden that can predispose toward the appearance of cancer. Interestingly, tissue, age and even gender differences in the robustness of the circadian cycle have been documented ^52^, something that might be of importance in cancer evolution. Our analysis using TCGA data and mouse xenografts suggest this idea of higher chances of tumour development and a faster evolution when CRY1 protein levels are reduced. Furthermore, this should also be considered for cancer treatment, as cells depleted for CRY1 are more resistant to DSB-inducing agents, and the same tendency is suggested from the xenografts models. The exploitation of the connection between circadian rhythms and cancer treatment efficiency is known as chronotherapy ^53^, and there is discussion if this might be applied to radiotherapy ^54^. We have even observed this effect in a real-life clinical setup, albeit with a modest impact and, interestingly, only on specific tumour samples. The reasons behind those differences and, specifically, if this reflects the loss of CRY1 regulation in specific tumours must be explored in the future to understand the therapeutic potential of chronoradiotherapy.

## Methods

### Cell culture and cell lines

U2OS cells and MEFs were grown in high glucose DMEM (Sigma-Aldrich) supplemented with 10% foetal bovine serum (Sigma-Aldrich), 2 mM L-glutamine (Sigma-Aldrich), 100 units/ml penicillin, and 100 μg/ml streptomycin (Sigma-Aldrich). U2OS cells stably expressing GFP-CtIP, GFP-CCAR2, Cherry-CtIP and Cherry-MDC1 were grown in a standard medium supplemented with 0.5 mg/mL G418 (Gibco, 15140122). U2OS cells harbouring a single copy of the reporter constructs DR-GFP, SA-GFP and EJ5-GFP were grown in a standard medium supplemented with 1 μg/mL puromycin (Sigma, P8833).

RPE1 cells were grown in high glucose DMEM/F-12 medium (Sigma-Aldrich) supplemented with 10% foetal bovine serum (Sigma-Aldrich), 2 mM L-glutamine (Sigma-Aldrich), 100 units/ml penicillin, and 100 μg/ml streptomycin (Sigma-Aldrich). RPE1 cells stably expressing shRNAs against CtIP, CCAR2 and a control sequence (shScr) were grown in this medium supplemented with 5μg/mL puromycin (Sigma, P8833).

### Mice

6-8 week-old *Crl:NU(NCr)-Foxn1^nu^* mice (nude mice) for xenograft studies were purchased from Charles River laboratories. The xenograft experiments were conducted under a protocol approved by the CABIMER bioethics committee and the Junta de Andalucía (Protocol 26/02/2021/025). All mice were housed in a pathogen-free environment at CABIMER animal facility and were handled in strict accordance with the Spanish and European Animal experimentation regulations’’

### Human

Patient mutational datasets were retrieved from the International Cancer Genome Consortium (ICGC) Data Portal TCGA Breast Cancer (BRCA) datasets. CRY1 and CCAR2 expression levels, survival and radiation therapy information from each donor were obtained using the UCSC Xena web tool.

Survival data was obtained from the Radiotherapy Service at the University Hospital Virgen Macarena, Seville Spain, from all patients treated with radiotherapy as a primary form of treatment between May 2018 to October 2023. In all cases, in addition to the type of cancer, the gender and age (Suppl. Table 5), the date of treatment initiation, the date of the last known interaction (either the death of the patient or the last known follow up), and approximate time of irradiation was recorded. Each patient was always treated at the same time of the day. Patients were split in two cohorts, according to the time of irradiation, those treated in the first half of the day (8:00 to 14:00; morning) or the second half (14:00 to 20:00).

### siRNAs, plasmids, and transfections

siRNA duplexes were obtained from Sigma-Aldrich or Dharmacon (Suppl. Table 2) and were reversely transfected using RNAiMax Lipofectamine Reagent Mix (Life Technologies), according to the manufacturer’s instructions. Plasmid transfection was carried out using FuGENE 6 Transfection Reagent (Promega) according to the manufacturer’s protocol.

### Circadian synchronization experiments

Circadian synchronization was performed by treating cells with either 1 μM of Dexamethasone (Sigma), or 10 μM of Forskolin (Sigma) or mock treated with EtOH or DMSO as a control, respectively, for 2 hours. Release from synchronization was performed by exchanging the medium with fresh one.

### Immunoprecipitation

U2OS cells were harvested in lysis buffer (50 mM Tris–HCl, pH 7.5, 50mM NaCl, 1 mM EDTA, 0.2 % Triton X-100, 1X protease inhibitors (Roche), 1X phosphatase inhibitor cocktail 1 (Sigma)) and incubated for 30 min on ice with Benzonase (100 U/mL). Protein extracts (500 μg-1 mg) were then subjected to preclearing with 20 μL of either magnetic protein G (for Rabbit antibodies) or Protein A (for Mouse antibodies) Dynabeads (Invitrogen) for 1 hour at 4°C. Then, precleared extracts were incubated overnight with 0.5 μg of CRY1 antibody (Rabbit, A302-614A), 1 μg of CCAR2 antibody (Rabbit, A300-433A-1), 1.5 μL of CCAR2 antibody (Mouse, 5857) or the equivalent amount of IgG (Mouse or Rabbit) as the negative control. Subsequently, extracts were incubated with the corresponding magnetic protein beads and after that they were washed three times with lysis buffer with 0.02% of Triton X-100. The precipitate was eluted in 30 μl of Laemmli buffer 2X. The antibodies used for immunoprecipitation can be found on Suppl. Table 3.

### Immunofluorescence and microscopy

All the experiments were performed in close collaboration with the microscopy unit at CABIMER: For foci visualization, cells were seeded on 12 mm coverslips, and after the different treatments, cell were washed once with PBS before following the immunofluorescence procedure.

For RPA and BRCA1 foci analysis, cells were pre-extracted with Pre-extraction Buffer (25 mM Tris–HCl, pH 7.5, 50 mM NaCl, 1 mM EDTA, 3 mM MgCl_2_, 300 mM sucrose, and 0.2% Triton X-100) for 5 min on ice. Then, cells were fixed on ice with 4% paraformaldehyde (w/v) in PBS for 20 min. For RIF1, 53BP1 and γH2AX foci analysis, cells were treated for 20 min on ice with 4% paraformaldehyde (w/v) in PBS, and after two washes with PBS 1X, they were permeabilized with 0.25% Triton X-100 in PBS. For RAD51 foci analysis, cells were pre-extracted with 0.1% Triton X-100 in PBS for 5 min on ice. Then, cells were fixed on ice with 4% paraformaldehyde (w/v) in PBS for 20 min on ice.

In all cases, following two washes with PBS, cells were blocked for 1 h with 1% BSA in PBS, co-stained with the appropriate primary antibodies (Suppl. Table 3) in blocking solution overnight at 4 °C, washed again with PBS and then co-immunostained with the appropriate secondary antibodies (Suppl. Table 4) in blocking buffer. After washing with PBS and dehydration with 100% ethanol, coverslips were mounted into glass slides using Vectashield mounting medium with DAPI (Vector Laboratories). In the case of QIBC experiments, cells were stained with 0.5 μg/mL DAPI in PBS for 5 min prior to drying with ethanol and were mounted using Vectashield mounting medium without DAPI (Vector Laboratories). Images were acquired and analysed using a Leica fluorescence microscope. The analysis of the number of foci formation in all cases was performed automatically using FIJI software and a FIJI custom macro that uses the bult-in Find Maxima function in FIJI, allowing the determination of discrete local maximum intesities inside the nucleus, being these discrete maximum spots regarded as foci.

### Single-molecule analysis of resection tracks

To measure the length of the resected DNA fragments SMART (single-molecule analysis of resection tracks) assay was performed or directly lysing and stretching the DNA on coverslip (SMART-tilt) (Altieri et al., 2020). U2OS cells were grown in the presence of 32 µM CldU (Sigma, C6891) for 24 h. They were then irradiated (10 Gy) for 1 h at 37°C and harvested using Trypsin. Cells were centrifuged at 500 g for 3 min at 4 °C, resuspended in PBS and mixed in a 1:8 proportion with unlabelled cells. For DNA stretching, 2.5 µl of this mix were lysed using a spreading buffer (50 mM EDTA, 0.5% SDS, 200 mM Tris-HCl pH 7.4) to later stretch the nucleic acid fibers on silanized slides (Sigma, S4651), tilting the slides to a 15°. After air-drying the DNA fibers for 10 min, they were fixed in 3:1 methanol/acetic acid at −20 °C for 15 min. Slides were then washed twice in PBS, incubated in 70% ethanol overnight at 4°C and washed twice again in PBS. For immunodetection, samples were blocked with 5% BSA (Sigma, A4503) in PBS for 30 min at room temperature (RT) and incubated with anti-BrdU/CldU (Abcam, ab6326) diluted in blocking buffer for 1 h at 37°C. Slides were washed with PBS twice and incubated with the fluorescent secondary antibody (Alexa Fluor 488 anti-rat, Molecular Probes A-11006) diluted in blocking buffer for 1 h at RT. Finally, samples were washed twice in PBS, dried and mounted with ProLong Gold Antifade Reagent (Molecular Probes, P36930). DNA fibers images were acquired with a Leica Thunder Microscope with automatized stage and a 63X oil immersion objective. For each experiment, at least 100 DNA fibers were measured using the Fiji (ImageJ) image software analysis.

### SDS-PAGE and western blot analysis

Protein extracts were prepared in 2X Laemmli buffer (4% SDS, 20% glycerol, 125 mM Tris–HCl, pH 6.8) and boiled at 100 °C for 5 min. Then, samples were resolved by SDS–PAGE and transferred to Amersham™ Protran™ Premium NC nitrocellulose membranes (Cytiva Life Sciences). Membranes were blocked with 5% Skim Milk (Difco) in 0.1% Tween-20, 0.01% Sodium Azide supplemented 1X TBS. After that, membranes were blotted with the appropriate primary antibody (Suppl. Table 3) and infrared dyed secondary antibodies (Suppl. Table 4) diluted in Odyssey Blocking Buffer (LI-COR) supplemented with 0.1% Tween-20 and 0.01% Sodium Azide. Membranes were air-dried in the dark and scanned in an Odyssey Infrared Imaging System (LI-COR), and images were analysed with Image Studio software (LI-COR).

### Cell cycle analysis

Cells were fixed with cold 70% ethanol for at least 2 hours, incubated with 250 μg/ml RNase A (Sigma Aldrich) and 10 μg/ml propidium iodide (Sigma Aldrich) at 37 °C for 30 min and analysed with a LSRFortessa™ Cell Analyzer (BD) Flow Cytometer. Cell cycle distribution data were further analysed using ModFit LT 5.0 software (Verity Software House Inc).

### HDR and NHEJ analysis

U2OS cells bearing a single copy integration of the reporters DR-GFP (Gene conversion), SA-GFP (SSA) or EJ5-GFP (NHEJ) were used to analyse the different DSB repair pathways. In all cases, 60.000 cells were plated in 6-well plates in duplicate. One day after seeding, cells were transfected with the indicated siRNA or plasmids. The next day, each duplicate culture was infected with lentiviral particles containing I-SceI– BFP expression construct at MOI 10 using 8 µg/ml polybrene in 2 ml of DMEM. Then, cells were left to grow for an additional 24 h before changing the medium for fresh DMEM. 48 h after viral transduction, cells were washed with PBS, trypsinized, neutralized with DMEM, centrifuged for 3 min at 600 g, and collected by centrifugation and resuspended in 300 µl of PBS. Samples were analysed with a LSRFortessa™ Cell Analyzer (BD) Flow Cytometer.

### UV laser micro-irradiation

For the analysis of laser-localized fusion proteins, cells were seeded on μ-slide glass bottom plates (Ibidi). Experiments were imaged with an ORCA Flash 4.0 sCMOS camera (Hamamatsu) using a Yokogawa CSU-W1 Confocal Scanner Unit (Gataca Systems) in an Axio Observer 7 microscope (Zeiss) at 37 °C and 5% CO_2_ conditions. Laser microirradiation was performed using a laser output of 20% power of a 355nm laser line (10mW; 19kHz, iLAS 3 Gataca Systems) through a UV-transmitting 100-X oil immersion objective (Zeiss). Laser stripes were created at the cell nucleus with a pixel size of 1, and 2 iterations per stripe using MetaXpress software (Molecular Devices). After irradiation, GFP or Cherry tagged protein accumulation at laser tracks was recorded at the indicated times. Signal intensity was measured using Fiji (ImageJ) by subtracting pre-irradiation signal to each time-point and represented as arbitrary units (a.u.).

### Clonogenic cell survival assays

Clonogenic survival assays were carried out seeding 500 cells in 6-well plates triplicates 24 hours prior to treatment. DSBs were produced by ionizing radiation (IR) or by acute treatment with topoisomerase inhibitor etoposide (VP16; Sigma). For IR, cells were exposed to 2 or 4 Gy or mock treated. In the case of VP16 treatments, cells were incubated for 1 h with 5 or 10 μM of the drug or vehicle (DMSO) as control. After two washes with PBS, a fresh medium was added, and cells were incubated at 37 °C for 7– 14 days to allow colony formation. Afterward, cells were stained and visualized in the solution of 0.5% Crystal Violet (Merck) and 20% ethanol (Merck). Once the colonies were stained, this solution was removed, and plates were washed with distilled water. The portion of surviving colonies at each dose was calculated by dividing the average number of visible colonies in treated versus control dishes.

### Subcutaneous xenografts in mice

Xenografts were generated by injecting 8×10^6^ cells (HCT116) subcutaneously into flanks of female Athymic Nude (Crl:NU(NCr)-*Foxn1^nu^*) mice (Charles River Laboratories). Each animal was injected with cells depleted for CRY1 (right flank) or control cells (left flank). Tumour volume was estimated according to the formula: (Width^2^ × Length)/2. The treatment with etoposide (VP16) started when tumours reached 100-200 mm^3^. Mice were injected with etoposide (20mg/kg) or DMSO twice per week and tumour volume and mice weight were measured also twice per week for 20 days. At the end of the treatment, mice were sacrificed, and tumours were removed to weigh them and acquire images.

### ChIP-Seq

Chromatin Immunoprecipitation was performed as previously described (Jimeno-González et al., 2015). Briefly, DSBs were induced in AsiSI-ER U2OS cells (which stably express AsiSI-ER) in low-glucose Dulbecco’s Modified Eagle Medium (DMEM) + 10% charcoal-stripped FBS containing 300 nM 4-OHT for 4h. Then, cells were crosslinked with 1% formaldehyde for 10 minutes at 37°C. Crosslinking reaction was quenched with 125mM glycine for 5 minutes and cells were washed 3 times with PBS. Cell pellets were resuspended in 2.5 mL lysis buffer A (5 mM Pipes pH 8.0, 85 mM KCl, 0.5% NP40, 1mM PMSF) supplemented with protease inhibitors and incubated for 10 minutes on ice. After centrifugation, nuclear fraction was resuspended in 1 mL of lysis buffer B (50 mM Tris HCl pH 8.1, 1% SDS, 10 mM EDTA, 1 mM PMSF) supplemented with protease inhibitors. Chromatin was sonicated using a Bioruptor Pico (Diagenode, B01060010), 5 cycles of 30 seconds “on” and 30 seconds “off” on ice-cold water. For each immunoprecipitation, 20-100 μg of chromatin and 4 μg of the indicated antibody was used in IP buffer (0,1% SDS, 1% TX-100, 2mM EDTA, 20 mM TrisHCl pH8, 150 mM NaCl) at 4°C o/n and then with 40 μL of pre-blocked (1 mg/ml BSA) Dynabeads protein A (ThermoFisher). Beads were sequentially washed with Wash buffer 1 (20 mM Tris HCl pH 8.1, 0.1% SDS, 1% Triton x-100, 2 mM EDTA, 150 mM NaC), Wash buffer 2 (20 mM Tris HCl pH 8.1, 0.1% SDS, 1% Triton x-100, 2 mM EDTA, 500 mM NaCl) Wash buffer 3 (10 mM Tris HCl pH 8.1, 1% NP-40, 1% NaDoc, 1 mM EDTA, 250 mM LiCl), and twice with TE-buffer (10 mM Tris-HCL pH8, 1 mM EDTA pH8). ChIPmentation was carried out as previously described(Schmidl et al., 2015), using Tn5A Enzyme provided by the Proteomic Service of CABD (Centro Andaluz de Biología del Desarrollo). Tagmented DNA was then eluted with 1% SDS in TE at 65°C for 10 minutes and crosslinking was reverted at 65°C overnight. Protein was degraded with Proteinase K for 2 hours at 37°C. DNA was purified using QIAGEN PCR purification Kit (QIAGEN, 28106). Libraries were amplified for N-1 cycles (being N the optimum Cq determined by qPCR reaction) using NEBNext High-Fidelity Polymerase (New England Biolabs, M0541). Libraries were purified with Sera-Mag Select Beads (GE Healthcare, 29343052) and sequenced using Illumina NovaSeq 6000 SP300 cycles 2×150 pb long (paired-end), XP protocol. All sequencing was performed at the Genomics Unit of CABIMER. Raw sequencing files (fastq) quality was assessed using FastQC 0.12.0 (https://www.bioinformatics.babraham.ac.uk/projects/fastqc/) and aligned to the reference human genome (version hg19) using bwa 0.7.17 (https://bio-bwa.sourceforge.net/). Mapped sequences were sorted and indexed using samtools 1.19.2 (http://www.htslib.org). Coverage of mapped reads in bigwig format was obtained using deepTools 3.5.3 bamCoverage function, setting normalization method to CPM. Bigwig files were visualized using Integrative Genomics Viewer 2.17.0. Average ChIP-seq plots of 214 experimentally characterized AsiSI sites(Iannelli et al., 2017) were generated using deepTools computeMatrix and plotProfile functions. Sequencing was performed at the Genomics Unit of CABIMER:

### SBS3 mutational signature analysis

Association analysis of homologous recombination deficiency signature (Cosmic SBS3) counts to CRY1 expression was performed as follows. RNA-seq data from patient cohorts was retrieved using the UCSC Xena web tool (https://xenabrowser.net/). Patient stratification into high- or low-expressing subclasses was performed by subdividing expression data into quartiles of CRY1 expression and utilizing the first quartile as low-expression samples and the fourth quartile as high-expression samples. SBS3 mutational signature per sample was obtained using SigProfilerMatrixGenerator (Islam et al., 2020). and retrieving SBS3 mutational signature per CRY1 group.

### Neutral comet assays

U2OS cells were treated with ionizing radiation (10 Gy) and incubated for 24 hours or mock treated. After trypsinization and neutralization, cells were resuspended in PBS at a concentration of 500.000 cells per mL and mixed in a 1:1 ratio with low-melting agarose at 37°C. 20 uL of cell suspension was dropped onto agarose covered microscope slides and, after drying, slides were incubated in lysis buffer (Trevigen) overnight at 4°C. After lysis, electrophoresis of the DNA trapped in the agarose was performed at 1 V/cm for 45 min in TEB 1X buffer. DNA was stained with SYBR gold for 10 min and slides were washed in water and further dried at 37°C. At least 200 cells were analyzed using a Leica DMI8 Thunder fluorescence microscope and tail length was analyzed using ImageJ.

### Gross chromosomal aberration (GCA) detection in mitotic spreads

Gross chromosomal aberrations were analyzed on DAPI-stained mitotic spreads. U2OS cells treated as described in figure legends were irradiated with a total dose of 2 Gy and allowed to recover for 7 hours. Within these 7 hours, cells were treated with 2 mM caffeine (Sigma, C0750) for the last 5 h to allow cells with gross chromosomal aberrations (GCAs) to enter mitosis bypassing the ATM/ATR G2/M checkpoint (REF). For the last 3 hours, they were treated with 0.1 μg/ml Demecolcine (Sigma, D9125) to induce chromosome condensation. Cells were then harvested and treated with 8 g/L sodium citrate at 37°C for 12 min to produce the hypotonic shock of the cells. Afterwards, cells were pre-fixed by adding 1 mL of Fixation Solution (a mixture of methanol and acetic acid in a 3:1 ratio), washed once, and fixed in Fixation Solution. After fixation, cells were resuspended in the fixative and cell suspension was dropped onto glass microscope slides (previously stored at −20 °C to allow the spread of the metaphases). Then, slides were air-dried and mounted with Vectashield Antifade mounting medium containing DAPI (Vector Laboratories, H-1200). Samples were analyzed using a Leica DMI8 Thunder fluorescence microscope and the number of GCAs per mitosis was scored in 50 cells per experiment using ImageJ.

### QUANTIFICATION AND STATISTICAL ANALYSIS

Statistical significance was determined with the test indicated in the corresponding figure legend using PRISM software (Graphpad Software). Statistically significant differences were labeled with one, two, three or four asterisks if P < 0.05, P < 0.01, P < 0.001 or P < 0.0001, respectively. Specific replicate numbers (n) for each experiment can be found in the corresponding figure legends.

## DATA AVAILABILITY

All relevant data are included in the manuscript. Raw data will be provided upon request. Source data are provided with this paper. ChIP-seq data are deposited at GEO under the accession number GSE261427.

## ACKNOWLEDGEMENTS.

This work was funded by the R+D+I grants PID2019-104195G and PID2022-136791NB-I00 funded by MICIU/AEI/10.13039/501100011033/ FEDER/UE, the grant P18-RT-1204 from the Consejería de Transformación Económica, Industria, Conocimiento y Universidades, Junta de Andalucía and the grant PRYGN246626HUER from the Asociación Española Contra el Cáncer. The mouse work in ALC laboratory was funded by the R+D+I grant PID2020-119329RB-I00, SJ lab is funded by the grant PID2022-140603NB-I00 and SJG lab is funded by the grant PID2022-139253N, all three funded by MICIU/AEI/10.13039/501100011033/ FEDER/UE. MC-P is the recipient of a MSCA-IF-2020 (coDNAres) that cover his salary and part of the reagents. AR-F is funded with a FPU fellowship from the Spanish Ministry of Education and PA was supported by a postdoctoral fellowship from the Junta de Andalucía and the European Social Fund (202099908313486), and by a postdoctoral grant from the Scientific Foundation of the Spanish Association Against Cancer (POSTD234422AGUI). CABIMER is supported by the regional government of Andalucía (Junta de Andalucía). We thank José Carlos Reyes-Rosa for critical reading of the manuscript. We also want to thank the Microscopy and Genomic Unit of CABIMER for their support in the experiments.

## AUTHORS CONTRIBUTIONS

PH and CCR conceived the project. PH and ARF designed the experiments. ARF, CCR, SJ, MCP, PA, SJG performed experiments. AW and HM collected patient data. AJLC supervised the mouse work. PH wrote the manuscript with input from all the authors.

## ETHICS&INCLUSION

All authors are based on Seville (Spain) as a collaboration of the Research Centre CABIMER and the local University Hospital Virgen Macarena. Research using patient clinical data was approved by the Ethical Board of the Hospital. The animal experimentation protocol was approved by the CABIMER Ethics Committee for Animal Experimentation (CEEA) and the Junta de Andalucía (project 26/02/2021/025) and the experiments were performed in compliance with the Spanish and European regulations.

