## Supplementary figures and images for "Circadian regulation of Homologous Recombination by Cryptochrome1-mediated dampening of DNA end resection"

### Supplemental figure 1

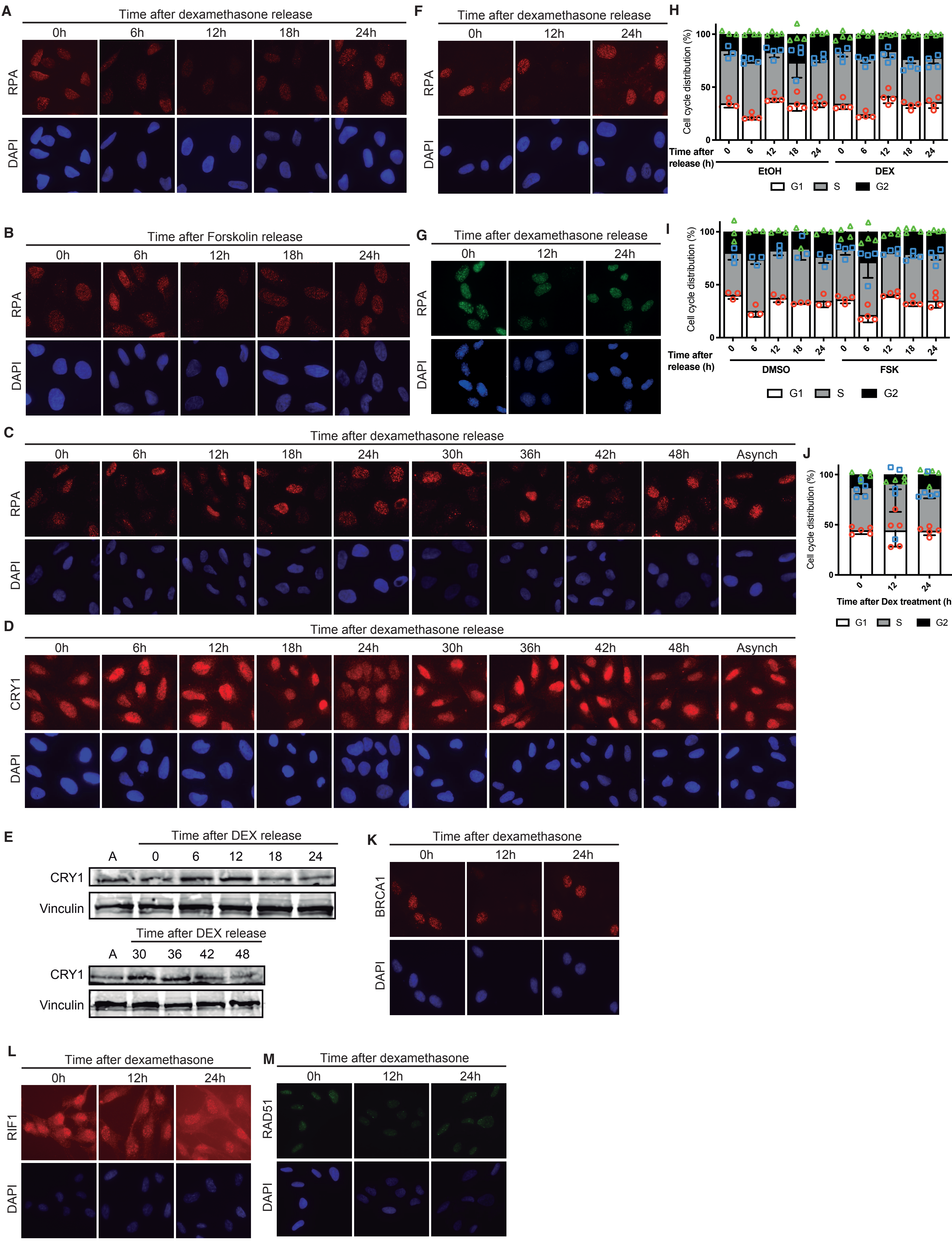


Romero-Franco et.al. Sup Figure 1

### Supplemental figure 4

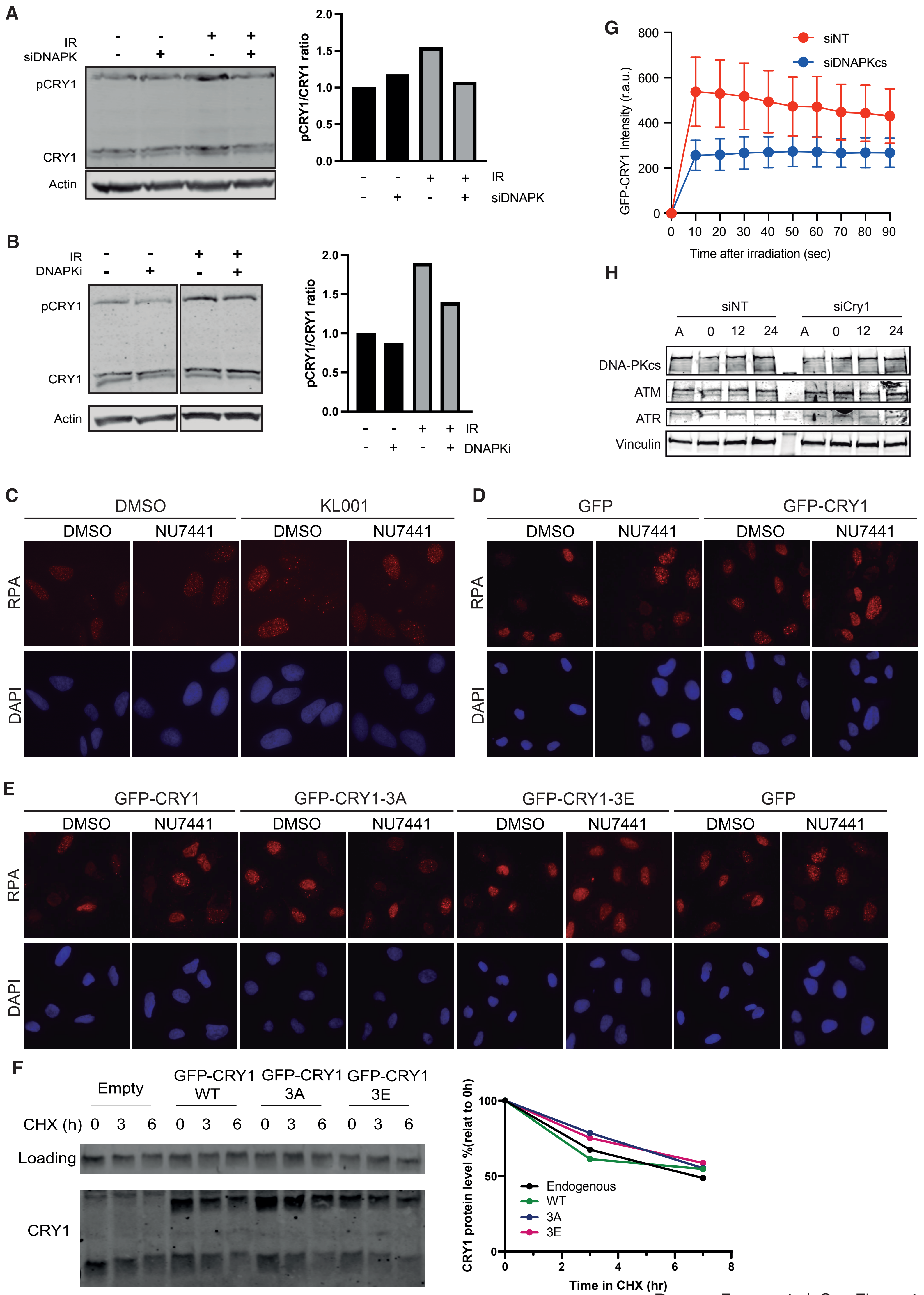


Romero-Franco et.al. Sup Figure 4

### Supplemental figure 5

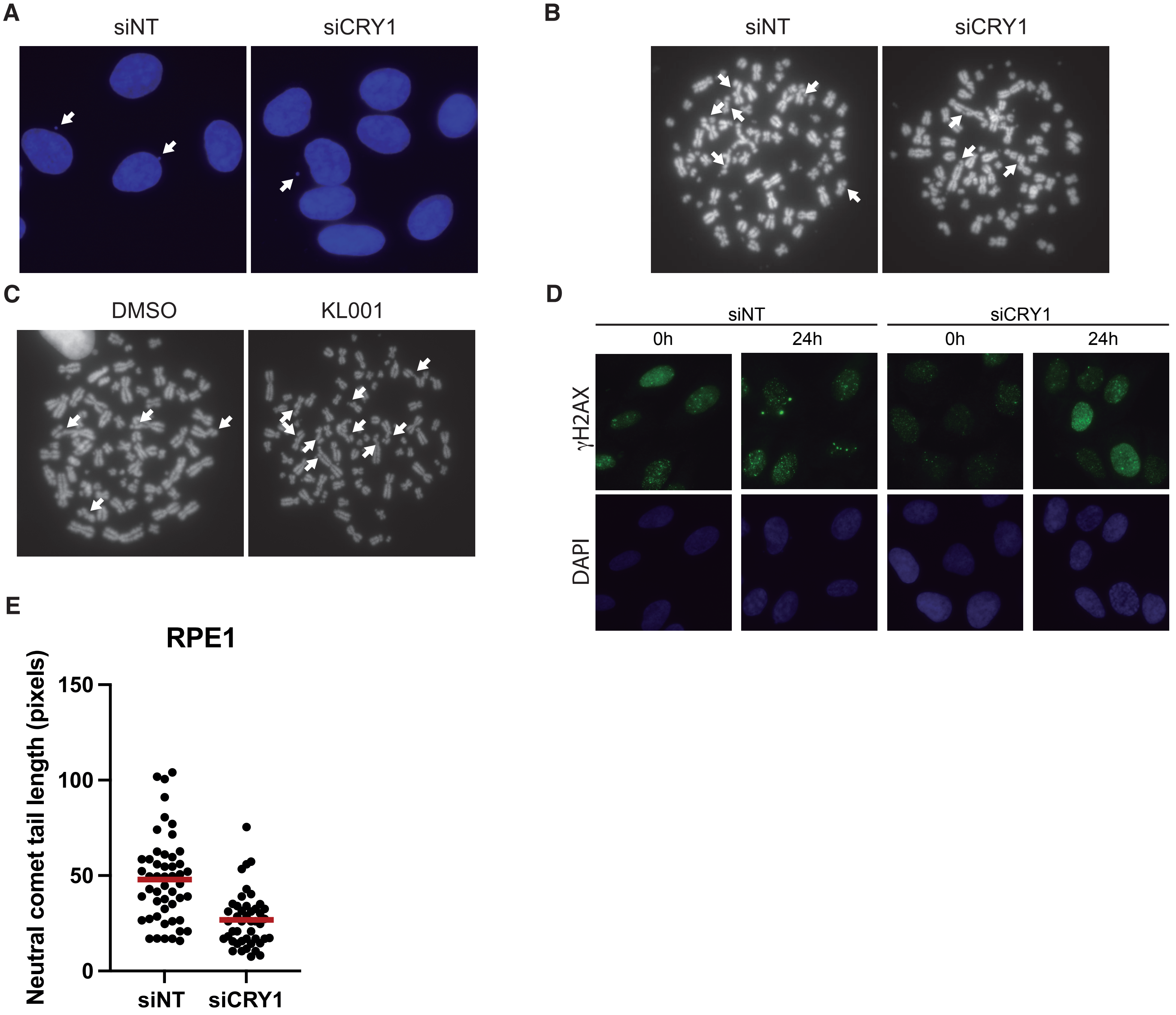


Romero-Franco et.al. Sup Figure 5

### Supplemental figure 6

A

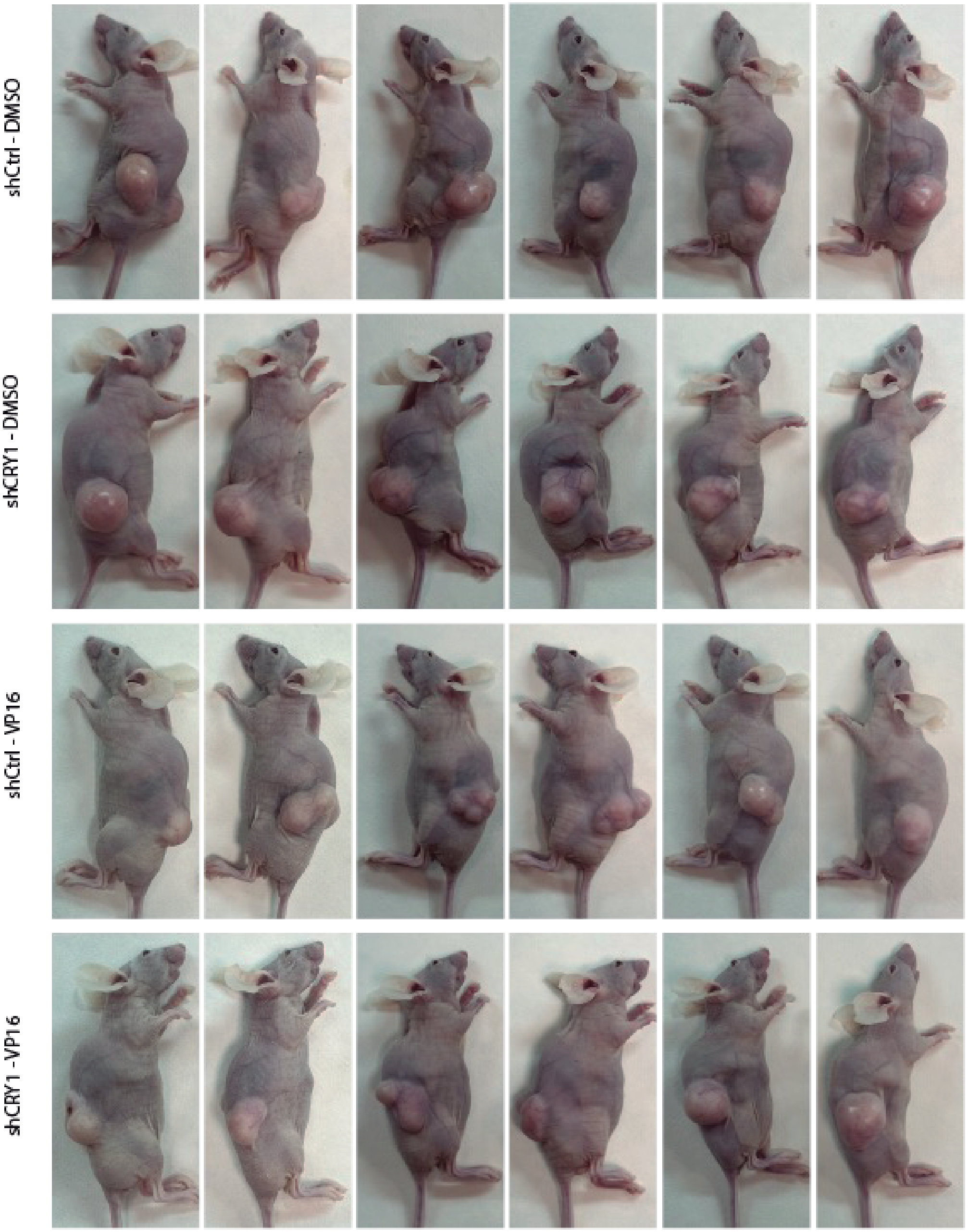


B

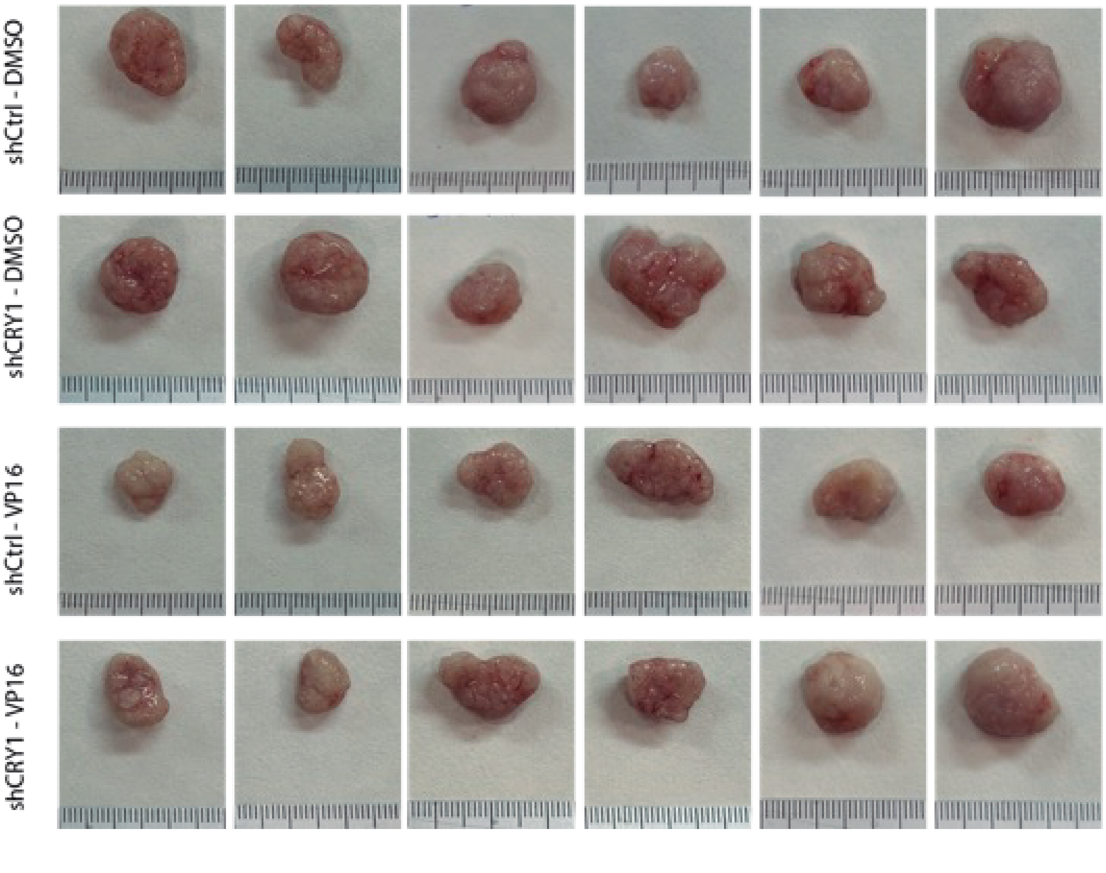


Romero-Franco et.al. Sup Figure 6
