## Supplemental figure 2 for "Circadian regulation of Homologous Recombination by Cryptochrome1-mediated dampening of DNA end resection"

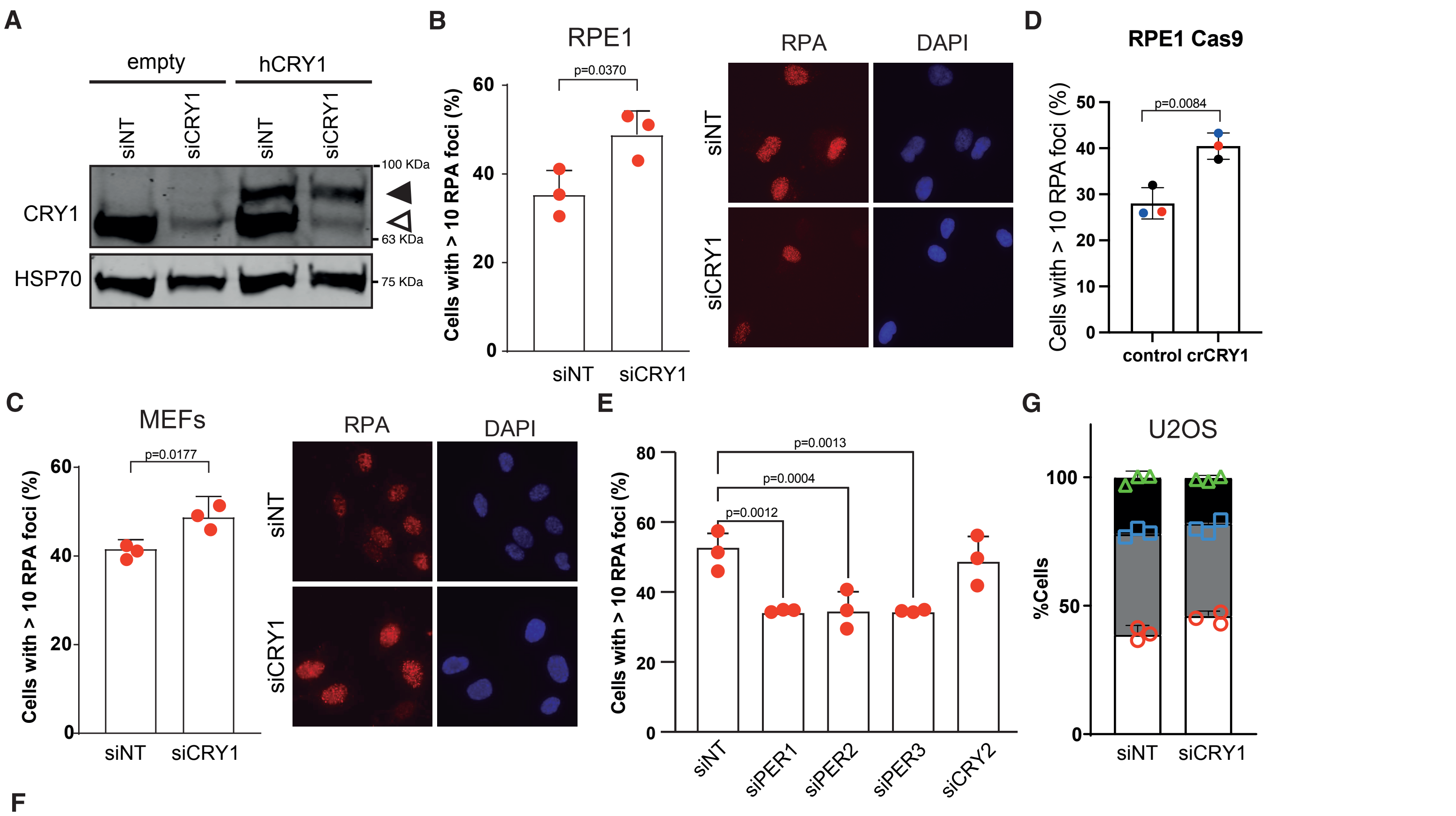


F

| | | siNT 0 | siNT 4 | siNT 8 | siNT 12 | siNT 16 | siNT 20 | siNT 24 | siCRY1 0 | siCRY1 4 | siCRY1 8 | siCRY1 12 | siCRY1 16 | siCRY1 20 | siCRY1 24 |
|---|---|---|---|---|---|---|---|---|---|---|---|---|---|---|---|
| siNT | 0 | nd | ns | ns | 0.04 | ns | ns | ns | ns | ns | ns | ns | ns | ns | ns |
| | 4 | | nd | ns | ns | ns | ns | ns | ns | ns | ns | ns | ns | ns | ns |
| | 8 | | | nd | ns | ns | ns | ns | 0.03 | <0.01 | <0.01 | 0.03 | ns | 0.01 | 0.02 |
| | 12 | | | | nd | ns | ns | <0.01 | <0.01 | <0.01 | <0.01 | <0.01 | <0.01 | <0.01 | <0.01 |
| | 16 | | | | | nd | ns | ns | <0.01 | <0.01 | <0.01 | 0.03 | ns | 0.01 | 0.02 |
| | 20 | | | | | | nd | ns | ns | ns | ns | ns | ns | ns | ns |
| | 24 | | | | | | | nd | ns | ns | ns | ns | ns | ns | ns |
| siCRY1 | 0 | | | | | | | | nd | ns | ns | ns | ns | ns | ns |
| | 4 | | | | | | | | | nd | ns | ns | ns | ns | ns |
| | 8 | | | | | | | | | | nd | ns | ns | ns | ns |
| | 12 | | | | | | | | | | | nd | ns | ns | ns |
| | 16 | | | | | | | | | | | | nd | ns | ns |
| | 20 | | | | | | | | | | | | | nd | ns |
| | 24 | | | | | | | | | | | | | | nd |

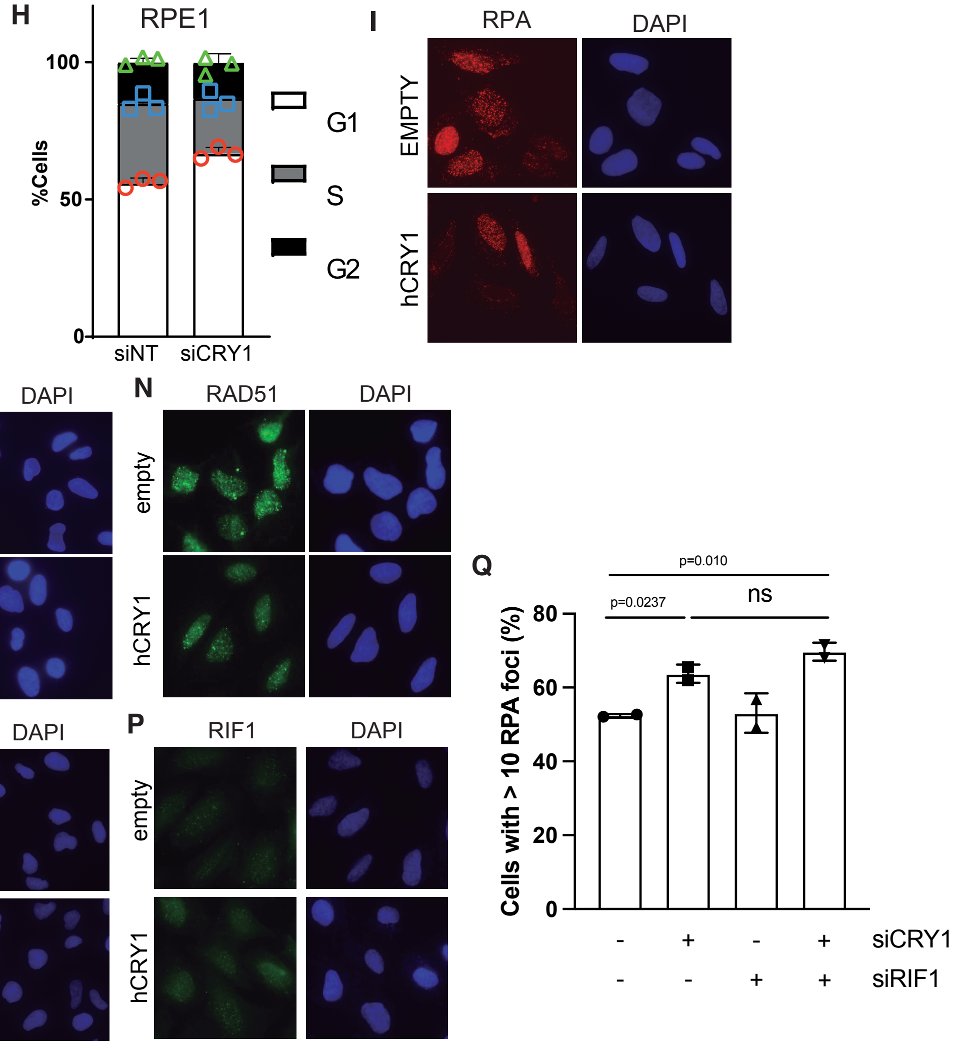


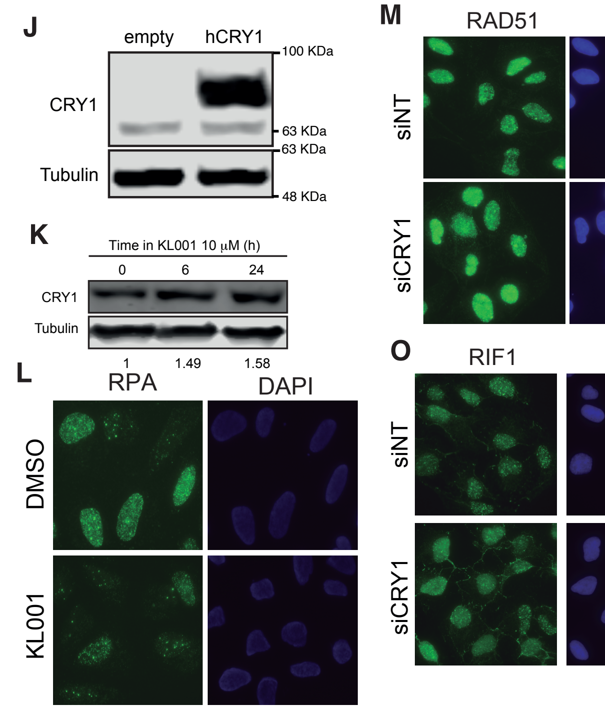


Romero-Franco et.al. Sup Figure 2
