## Supplemental figure 3 for "Circadian regulation of Homologous Recombination by Cryptochrome1-mediated dampening of DNA end resection"

A

| | | siNT 0 | siNT 4 | siNT 8 | siNT 12 | siNT 16 | siNT 20 | siNT 24 | siCtIP 0 | siCtIP 4 | siCtIP 8 | siCtIP 12 | siCtIP 16 | siCtIP 20 | siCtIP 24 |
|---|---|---|---|---|---|---|---|---|---|---|---|---|---|---|---|
| siNT | 0 | nd | ns | ns | 0.03 | ns | ns | ns | <0.01 | 0.01 | <0.01 | <0.01 | <0.01 | <0.01 | <0.01 |
| | 4 | | nd | ns | ns | ns | ns | ns | 0.01 | ns | 0.01 | ns | 0.01 | 0.02 | 0.02 |
| | 8 | | | nd | ns | ns | ns | ns | ns | ns | ns | ns | ns | ns | ns |
| | 12 | | | | nd | ns | ns | <0.01 | ns | ns | ns | ns | ns | ns | ns |
| | 16 | | | | | nd | ns | ns | ns | ns | ns | ns | ns | ns | ns |
| | 20 | | | | | | nd | ns | <0.01 | 0.03 | <0.01 | 0.03 | <0.01 | <0.01 | <0.01 |
| | 24 | | | | | | | nd | <0.01 | <0.01 | <0.01 | <0.01 | <0.01 | <0.01 | <0.01 |
| siCtIP | 0 | | | | | | | | nd | ns | ns | ns | ns | ns | ns |
| | 4 | | | | | | | | | nd | ns | ns | ns | ns | ns |
| | 8 | | | | | | | | | | nd | ns | ns | ns | ns |
| | 12 | | | | | | | | | | | nd | ns | ns | ns |
| | 16 | | | | | | | | | | | | nd | ns | ns |
| | 20 | | | | | | | | | | | | | nd | ns |
| | 24 | | | | | | | | | | | | | | nd |

B

| | | shNT 0 | shNT 12 | shNT 24 | shCTIP 0 | shCTIP 12 | shCTIP 24 | shCCAR2 0 | shCCAR2 12 | shCCAR2 24 |
|---|---|---|---|---|---|---|---|---|---|---|
| shNT | 0 | nd | 0.0264 | ns | 0.0033 | 0.0479 | 0.0236 | ns | ns | ns |
| | 12 | | nd | 0.0142 | ns | ns | ns | 0.0481 | 0.0147 | 0.0339 |
| | 24 | | | nd | 0.0002 | 0.0034 | 0.0173 | ns | ns | ns |
| shCtIP | 0 | | | | nd | ns | ns | 0.0009 | 0.0016 | 0.0003 |
| | 12 | | | | | nd | ns | 0.0213 | 0.0387 | 0.0082 |
| | 24 | | | | | | nd | 0.0482 | 0.0137 | 0.0374 |
| shCCAR2 | 0 | | | | | | | nd | ns | ns |
| | 12 | | | | | | | | nd | ns |
| | 24 | | | | | | | | | nd |

C

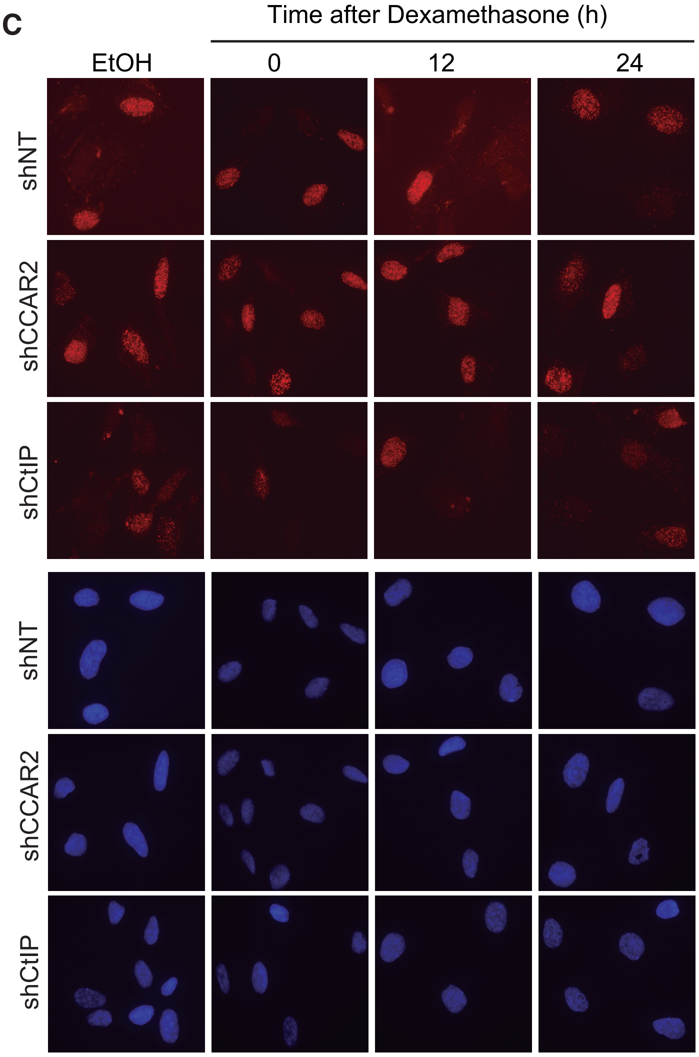


D

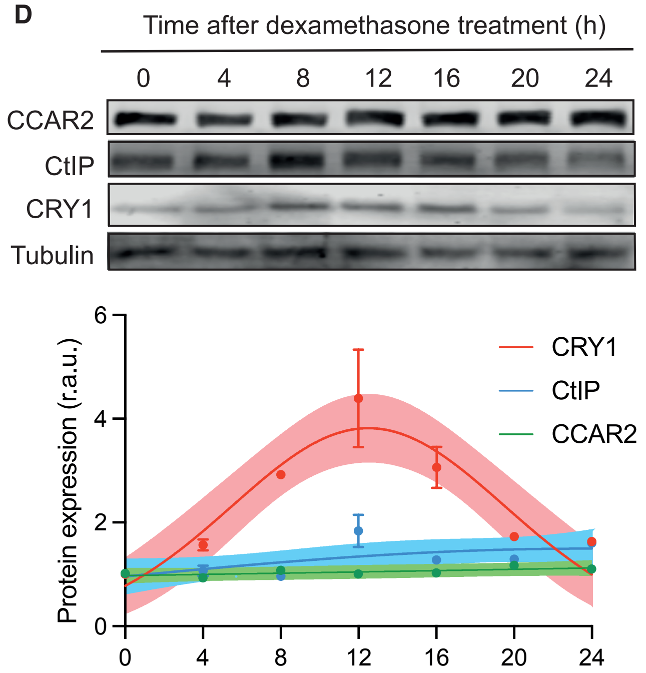


E

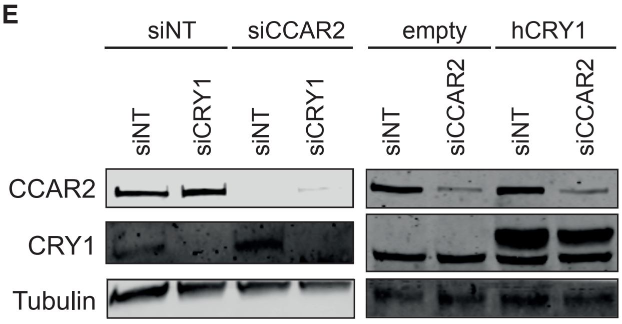


F

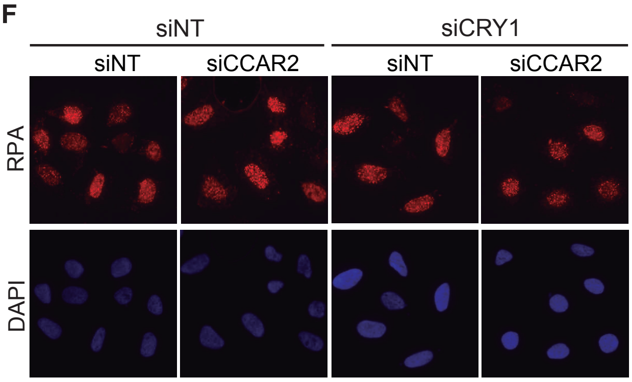


G

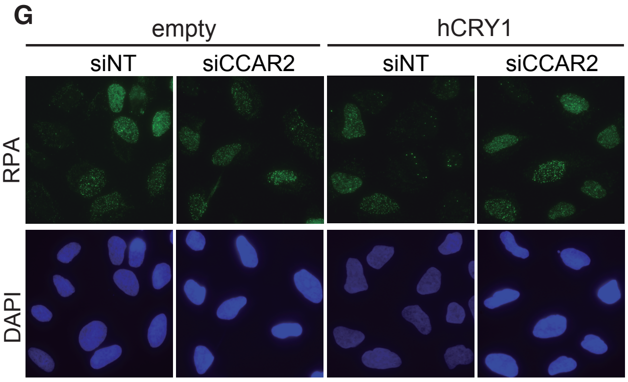


H

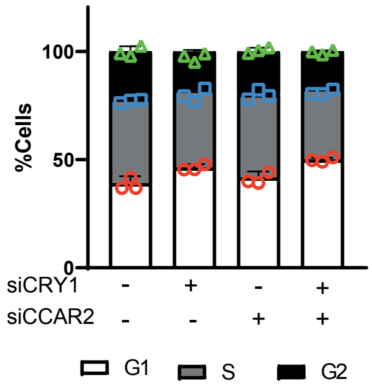


I

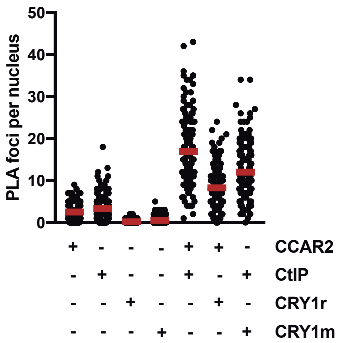


J

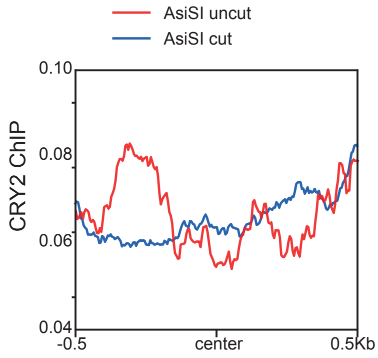
