## Supplemental information for "Circadian regulation of Homologous Recombination by Cryptochrome1-mediated dampening of DNA end resection"

**Supplementary Table 1: Statistical analysis of Figure 1D and 1E. Statistical significance between the denoted timepoints was calculated using an ANOVA test.**

| Timepoints (h) compared | p value for RPA recruitment (Figure 1D) | p value for CRY1 accumulation (Figure 1E) |
|---|---|---|
| 0 vs. 6 | 0.0346 | <0.0001 |
| 0 vs. 12 | 0.0198 | <0.0001 |
| 0 vs. 18 | 0.0393 | 0.6639 |
| 0 vs. 24 | 0.9999 | <0.0001 |
| 0 vs. 30 | 0.8861 | <0.0001 |
| 0 vs. 36 | 0.0288 | <0.0001 |
| 0 vs. 42 | 0.2078 | <0.0001 |
| 0 vs. 48 | 0.9654 | 0.0529 |
| 6 vs. 12 | 0.9999 | >0.9999 |
| 6 vs. 18 | 0.9999 | <0.0001 |
| 6 vs. 24 | 0.0264 | <0.0001 |
| 6 vs. 30 | 0.1888 | <0.0001 |
| 6 vs. 36 | >0.9999 | 0.9922 |
| 6 vs. 42 | 0.8561 | 0.9999 |
| 6 vs. 48 | 0.0099 | <0.0001 |
| 12 vs. 18 | 0.9994 | <0.0001 |
| 12 vs. 24 | 0.0153 | <0.0001 |
| 12 vs. 30 | 0.1062 | <0.0001 |
| 12 vs. 36 | >0.9999 | 0.9761 |
| 12 vs. 42 | 0.6325 | 0.9996 |
| 12 vs. 48 | 0.0060 | <0.0001 |
| 18 vs. 24 | 0.0300 | <0.0001 |
| 18 vs. 30 | 0.2145 | 0.0909 |
| 18 vs. 36 | >0.9999 | <0.0001 |
| 18 vs. 42 | 0.8953 | <0.0001 |
| 18 vs. 48 | 0.0112 | 0.9359 |
| 24 vs. 30 | 0.7932 | <0.0001 |
| 24 vs. 36 | 0.0221 | <0.0001 |
| 24 vs. 42 | 0.1580 | <0.0001 |
| 24 vs. 48 | 0.9910 | <0.0001 |
| 30 vs. 36 | 0.1569 | 0.0006 |
| 30 vs. 42 | 0.7930 | <0.0001 |
| 30 vs. 48 | 0.3802 | 0.8593 |
| 36 vs. 42 | 0.7904 | >0.9999 |
| 36 vs. 48 | 0.0084 | <0.0001 |
| 42 vs. 48 | 0.0550 | <0.0001 |

**Supplementary Table 2: siRNAs used in this work**

| Target gene | Description | Source/Reference/Sequence (5'-3') |
|---|---|---|

| | | |
|---|---|---|
| Control sequence | siNT | UGGUUUACAUGUCGACUAA, UGGUUUACAUGUUGUGUGA, UGGUUUACAUGUUUUCUGA, UGGUUUACAUGUUUUCCUA (mix) |
| CtIP | siCtIP | GCUAAAACAGGAACGAAUC |
| CRY1 3’UTR | siCRY1 3’UTR | ACAAAUUGAUAUUCUGAUAUA |
| CRY1 | siCRY1 | GAGGAUCUUGAUGCCAAUCUA |
| CCAR2 | siCCAR2 | GCUUAUAGUUCGAAGGUAC |
| CRY2 | siCRY2 | GAGGCCAUAGACAGAUCUAAA |
| PER1 | siPER1 | CCCGGACUCUCCACUGUUCAA |
| PER2 | siPER2 | AUAGGUGUUCAUCUAAGGUUA |
| PER3 | siPER3 | GCUAAAACAGGAACGAAUC |

**Supplementary Table 3:** Primary antibodies used in this study. WB, western blotting. IF, immunofluorescence. IP, Immunoprecipitation

| Primary antibody | Supplier | Reference | Application | Concentration |
|---|---|---|---|---|
| RPA32/RPA2 [9H8] (mouse) | Abcam | ab2175 | IF | 1:500 |
| Rat Anti-RPA32/RPA2 | Cell signaling | 2208 | IF | 1:500 |
| BRCA1 (D-9) (mouse) | Santa Cruz | sc-6954 | WB, IF | 1:500 |
| RAD51 (rabbit) | Abcam | ab133534 | IF | 1:500 |
| CENPF | Abcam | ab5 | IF | 1:500 |
| CCAR2 (DBC1) | Bethyl Laboratories | A300-433A-1 | WB | 1:1000 |
| Rabbit Anti-CCAR2 / DBC1 | Cell signaling | 5857 | WB | 1:1000 |
| Mouse Anti-pCCAR2/pDBC1 | Cell signaling | 4880 | IF | 1:100 |
| Rabbit Anti-CRY1 | Bethyl | A302-614A | IP, WB | 1:100, 1:2000 |

| | | | | |
|---|---|---|---|---|
| Mouse Anti-CRY1 | Santa Cruz | sc-1011006 | WB | 1:500 |
| Mouse Anti-CRY1 | Abcam | ab55649 | WB | 1:500 |
| Anti-IgG from mouse serum | Sigma | I8765 | IP | 1:100 |
| Anti-IgG from rabbit serum | Sigma | I8140 | IP | 1:100 |
| RIF1 (goat) | Santa Cruz | sc-55979 | IF | 1:100 |
| Rabbit Anti-RIF1 | Bethyl Laboratories | A300-569A | IF | 1:500 |
| γH2A.X (rabbit) | Cell Signaling | 2577L | IF | 1:500 |
| γH2A.X (mouse) | Abcam | ab22551 | IF | 1:500 |
| CtIP (rabbit) | Bethyl Laboratories | A300-487A | WB | 1:2000 |
| Rat Anti-BrdU | Abcam | ab6326 | IF | 1:100 |
| Rabbit Anti-CRY2 | Bethyl Laboratories | A302-615A | WB | 1:1000 |
| Rabbit Anti-53BP1 | Novus Biological | NB100-304 | IF | 1:500 |
| Rabbit Anti-β-Actin | Abcam | ab8227 | WB | 1:2000 |
| Mouse Anti-α-Tubulin | Sigma | T9026 | WB | 1:2000 |
| Mouse Anti-HSP70 | Santa Cruz | sc-24 | WB | 1:500 |

**Supplementary Table 4: Secondary antibodies used in this study.** WB, western blotting. IF, immunofluorescence.

| Secondary antibody | Supplier | Reference | Application | Concentration |
|---|---|---|---|---|
| Alexa Fluor 594 goat anti-mouse | Invitrogen | A11032 | IF | 1:1000 |
| Alexa Fluor 488 goat anti-rabbit | Invitrogen | A11034 | IF | 1:1000 |

| | | | | |
|---|---|---|---|---|
| Alexa Fluor 488 goat anti-rat | Invitrogen | A11006 | IF | 1:1000 |
| IRDye 680RD Goat anti-mouse IgG (H+L) | Li-cor | 926-68070 | WB | 1:5000-1:10000 |
| IRDye 800RD Goat anti-rabbit IgG (H+L) | Li-cor | 926-32211 | WB | 1:5000-1:10000 |
| IRDye 800CW Donkey anti-Goat IgG | Li-cor | 926-32214 | WB | 1:5000-1:10000 |

**Supplementary Table 5: Patient data.**

| **Cancer type** | **Number of Patients** | **% of Male patients** | **% Female patients** | **Average age of diagnosis** |
|---|---|---|---|---|
| **All** | 5751 | 45 | 55 | 64.1 |
| **Breast** | 2170 | 1 | 99 | 59.5 |
| **Glioma** | 102 | 60 | 40 | 57.7 |
| **Head and Neck** | 276 | 75 | 25 | 64.7 |
| **Lung** | 122 | 77 | 23 | 65 |
| **Melanoma** | 31 | 47 | 53 | 63.3 |
| **Prostate** | 1074 | 99.6 | 0.4 | 70.7 |

### SUPPLEMENTARY FIGURE LEGENDS

**Supplementary Figure 1. Resection oscillates following a circadian pattern. A**, Representative images of the experiment shown in Figure 1D. **B**, Representative images of the experiment shown in Figure 1E. **C**, Representative images of the experiment shown in Figure 1F. **D**, Representative images of the experiment shown in Figure 1G. **E**, U2OS cells exposed to dexamethasone for two hours were released by changing the medium. Protein samples were taken at the indicated timepoints, resolved in SDS-PAGE and blotted with the indicated antibodies. **F**, Representative images of the experiment shown in Figure 1H. **G**, Representative images of the experiment shown in Figure 1I. **H**, Cell cycle distribution of U2OS cells taken at the indicated times after the treatment with dexamethasone (left) or EtOH as a control (right) by FACs. **I**, Same as J but after treatment with forskolin or DMSO. **J**, Same as J but in RPE cells. **K**, Representative images of the experiment shown in Figure 1K. **L**, Representative images of the experiment shown in Figure 1L. **M**, Representative images of the experiment shown in Figure 1M.

**Supplementary Figure 2. CRY1 affects DNA end resection. A**, Representative western blot showing the depletion of CRY1 upon transfection with an siRNA and the expression of siRNA-resistant human CRY1 (hCRY1). Protein samples from U2OS cells transfected with a plasmid bearing a tagged form of CRY1 (hCRY1) or an empty vector and transfected with siRNA against CRY1 or a control sequence were resolved in SDS-PAGE and blotted with the indicated antibodies. Black triangle marks the ectopic version of CRY1 and white triangle the endogenous protein. **B**, Same as figure 2A but in RPE cells. **C**, Same as figure 2A but in MEFs cells. **D**, Same as B but in cells knockout for CRY1 using CRISPR. **E**, Same as figure 2A but in cells depleted for the indicated circadian factors. **F**, Statistical analysis of figure 2B using a 2-way ANOVA **G**, Cell cycle distribution of U2OS cells upon depletion or not of CRY1. **H**, Cell cycle distribution of RPE1 cells upon depletion or not of CRY1. **I**, Representative images of the experiment shown in Figure 2C. **J**, Representative western blot using samples from U2OS cells transfected with a plasmid bearing a tagged form of CRY1 (hCRY1) or an empty vector. **K**, Representative western blot using samples from U2OS cells treated with KL001 and taken at the indicated times. **L**, Representative images of the experiment shown in Figure 2D. **M**, Representative images of the experiment shown in Figure 2K. **N**, Representative images of the experiment shown in Figure 2L. **O**, Representative images of the experiment shown in Figure 2M. **P**, Representative images of the experiment shown in Figure 2N. **Q**, Same as panel B but in U2OS cells depleted of CRY1 and/or RIF1 as indicated.

**Supplementary Figure 3. CCAR2 and CRY1 cooperate controlling DNA end resection. A**, Statistical analysis of the data shown in figure 3A. nd: not determined. ns: not significant. Otherwise, the actual p value is shown. **B**, Statistical analysis of the data shown in figure 3B. Other details as in A. **C**, Representative images of the experiment shown in Figure 3B. **D**, U2OS ells exposed to dexamethasone for two hours were released by changing the medium. Protein samples were taken at the indicated timepoints, resolved in SDS-PAGE and blotted with the indicated antibodies. A representative image is shown on top. Protein kinetics was inferred using a nonlinear gaussian model from three replicates (bottom). 95% confidence intervals are marked as colored shades. **E**, Representative western blots using samples from U2OS cells transfected with the indicated siRNAs and bearing a plasmid expressing CRY1 or the empty vector, as depicted. Membranes were blotted with the mentioned antibodies. **F**, Representative images of the experiment shown in Figure 3C. **G**, Representative images of the experiment shown in Figure 3D. **H**, Cell cycle distribution of U2OS cells upon depletion of CRY1 and or CCAR2, as indicated. **I**, Proximity ligation assay using pairwise combinations of CCAR2, CtIP or CRY1 antibodies. CRY1r and CRY1m represent antibodies against CRY1 raised in rabbit or mouse, respectively. Incubations with a single antibody, as a negative control, are plotted. A representative experiment out of three with similar results is shown. **J**, Average recruitment of CRY2 at the 214 best AsiSI cutting sites measured by ChIP-seq in samples exposed (cut, blue line) or not to (uncut, red line) to tamoxifen to induce a cleavage at AsiSI sites.

**Supplementary Figure 4. CRY1 phosphorylation by DNA-PK affects resection. A**, Protein samples from U2OS transfected with an siRNA against DNAPKcs or a control sequence and exposed or not to IR, as indicated, were resolved in SDS-PAGE gels containing Phos-tag to separate the phosphorylated (pCRY1) and not phosphorylated (CRY1) form of CRY1. Membranes were blotted with the indicated antibodies. A representative western blot is shown on the left side, and quantification of the pCRY1/CRY1 ratio on the right side. **B**, same as A but in cells treated with a DNAPK inhibitor (DNAPKi) or the vehicle. **C**, Representative images of the experiment shown in Figure 4A**. D**, Representative images of the experiment shown in Figure 4B. **E**, Representative images of the experiment shown in Figure 4D. **F**, Cicloheximide (CHX) chase of CRY1 variants. Cells bearing the indicated construct were treated with CHX and protein samples taken at the indicated times. Proteins were resolved and bloated for CRY1. A representative western blot (left) and the quantification of CRY1 variants levels (right) is shown. **G**, Same as Figure 4G but in cells transfected with an siRNA against DNAPKcs or a control sequence. **H**, protein samples were taken at different time points after DEX release in both CRY1 depleted and control cells. Samples were resolved in SDS-Page and blotted using antibodies against the indicated PIKKs.

**Supplementary Figure 5. Genomic instability reacts to CRY1 levels. A**, Representative images of the experiment shown in Figure 5F**. B**, Representative images of the experiment shown in Figure 5G. **C**, Representative images of the experiment shown in Figure 5H. **D**, Representative images of the experiment shown in Figure 5H.

**Supplementary Figure 6. Effect of CRY1 levels in mouse Xenografts treated with etoposide. A**, Mice were grafted subcutaneously with HCT116 cells bearing a control shRNA on the left flank and harbouring an shRNA against CRY1 on the right side. 6 mice were treated with DMSO and 6 with etoposide (VP16), as described in the method section. Images of the tumour growth at the end of the experiment for each mice are shown. **B**, Photographs of the actual tumour after being extracted of the mice.

**Supplementary Movie 1: Recruitment of CRY1 at sites of DSBs.** U2OS cells harbouring GFP-CRY1 and Scartlet-MDC1 were laser microirradiated and imaged at different timepoints. Timestamps are included in the images. The recruitment of GFP-CRY1 is shown.

**Supplementary Movie 2: Recruitment of MDC1 at sites of DSBs.** U2OS cells harbouring GFP-CRY1 and Scartlet-MDC1 were laser microirradiated and imaged at different timepoints. Timestamps are included in the images. The recruitment of Scarlet-CRY1 is shown.

**Supplementary Movie 3: Recruitment of CtIP at sites of DSBs in cells bearing a control siRNA.** U2OS cells harbouring GFP-CtIP and transfected with a control siRNA were laser microirradiated and imaged at different timepoints. Timestamps are included in the images. The recruitment of GFP-CtIP is shown.

**Supplementary Movie 4: Recruitment of CtIP at sites of DSBs upon depletion of CRY1.** U2OS cells harbouring GFP-CtIP and transfected with a siRNA targeting CRY1 were laser microirradiated and imaged at different timepoints. Timestamps are included in the images. The recruitment of GFP-CtIP is shown.

**Supplementary Movie 5: Recruitment of CtIP at sites of DSBs in cells bearing an empty vector.** U2OS cells harbouring Cherry-CtIP and transfected with a vector harbouring GFP were laser microirradiated and imaged at different timepoints. Timestamps are included in the images. The recruitment of Cherry-CtIP is shown.

**Supplementary Movie 6: Recruitment of CtIP at sites of DSBs in cells upon overexpression of CRY1.** U2OS cells harbouring Cherry-CtIP and transfected with a vector harbouring GFP-CRY1 were laser microirradiated and imaged at different timepoints. Timestamps are included in the images. The recruitment of Cherry-CtIP is shown.

**Supplementary Movie 7: Early recruitment of wildtype CRY1 at sites of DSBs.** U2OS cells harbouring GFP-CRY1 were laser microirradiated and imaged at different timepoints. Timestamps are included in the images. The recruitment of GFP-CRY1 is shown.

**Supplementary Movie 8: Early recruitment of CRY1-3A at sites of DSBs.** U2OS cells harbouring GFP-CRY1-3A were laser microirradiated and imaged at different timepoints. Timestamps are included in the images. The recruitment of GFP-CRY1-3A is shown.

**Supplementary Movie 9: Early recruitment of CRY1-3E at sites of DSBs.** U2OS cells harbouring GFP-CRY1-3E were laser microirradiated and imaged at different timepoints. Timestamps are included in the images. The recruitment of GFP-CRY1-3E is shown.

**Supplementary Movie 10: Late recruitment of wildtype CRY1 at sites of DSBs.** Same as Supplementary Movie 7 but imaging at longer times.

**Supplementary Movie 11: Late recruitment of CRY1-3A at sites of DSBs.** Same as Supplementary Movie 8 but imaging at longer times.

**Supplementary Movie 12: Late recruitment of CRY1-3E at sites of DSBs.** Same as Supplementary Movie 9 but imaging at longer times.

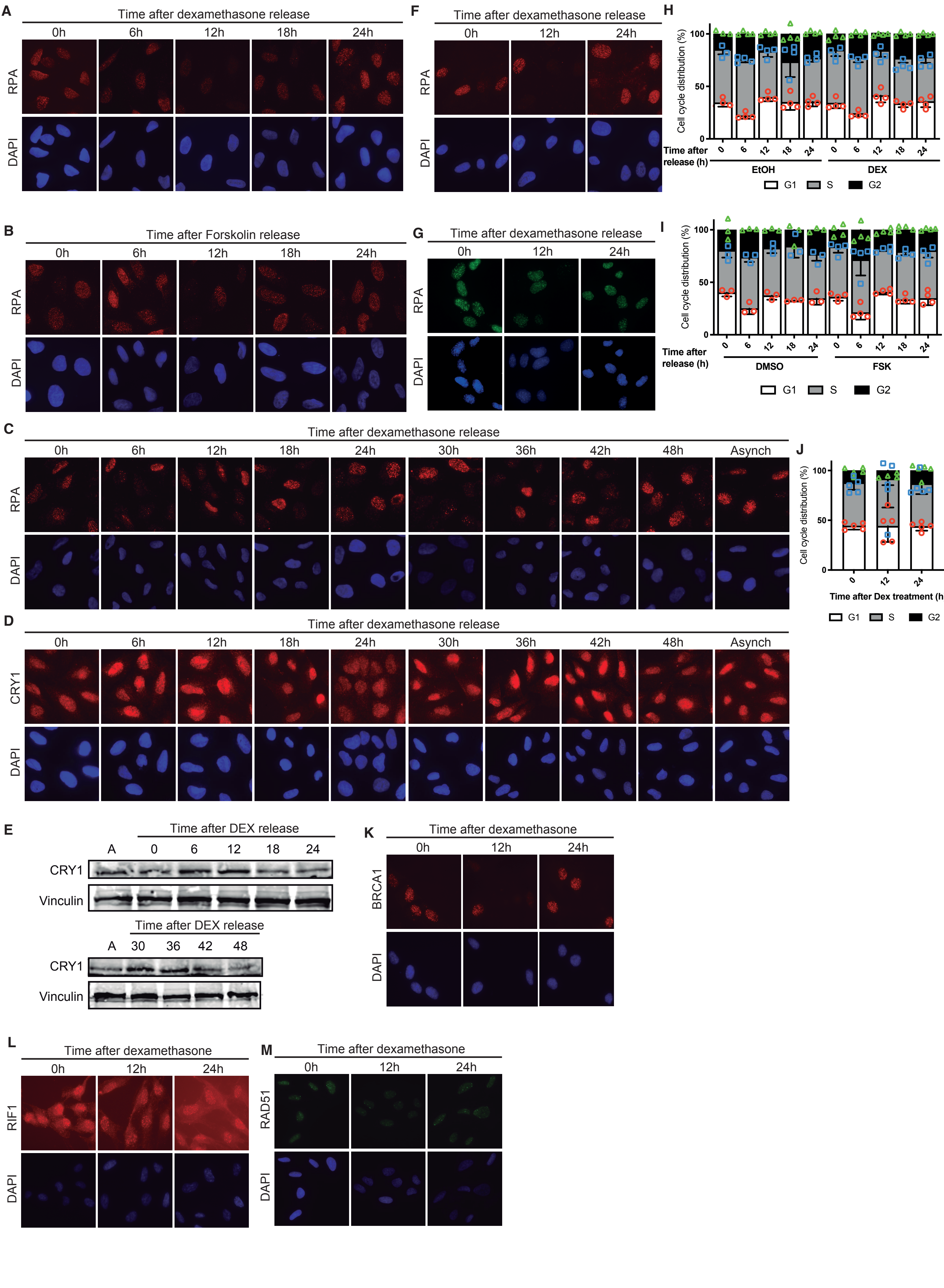


Romero-Franco et.al. Sup Figure 1

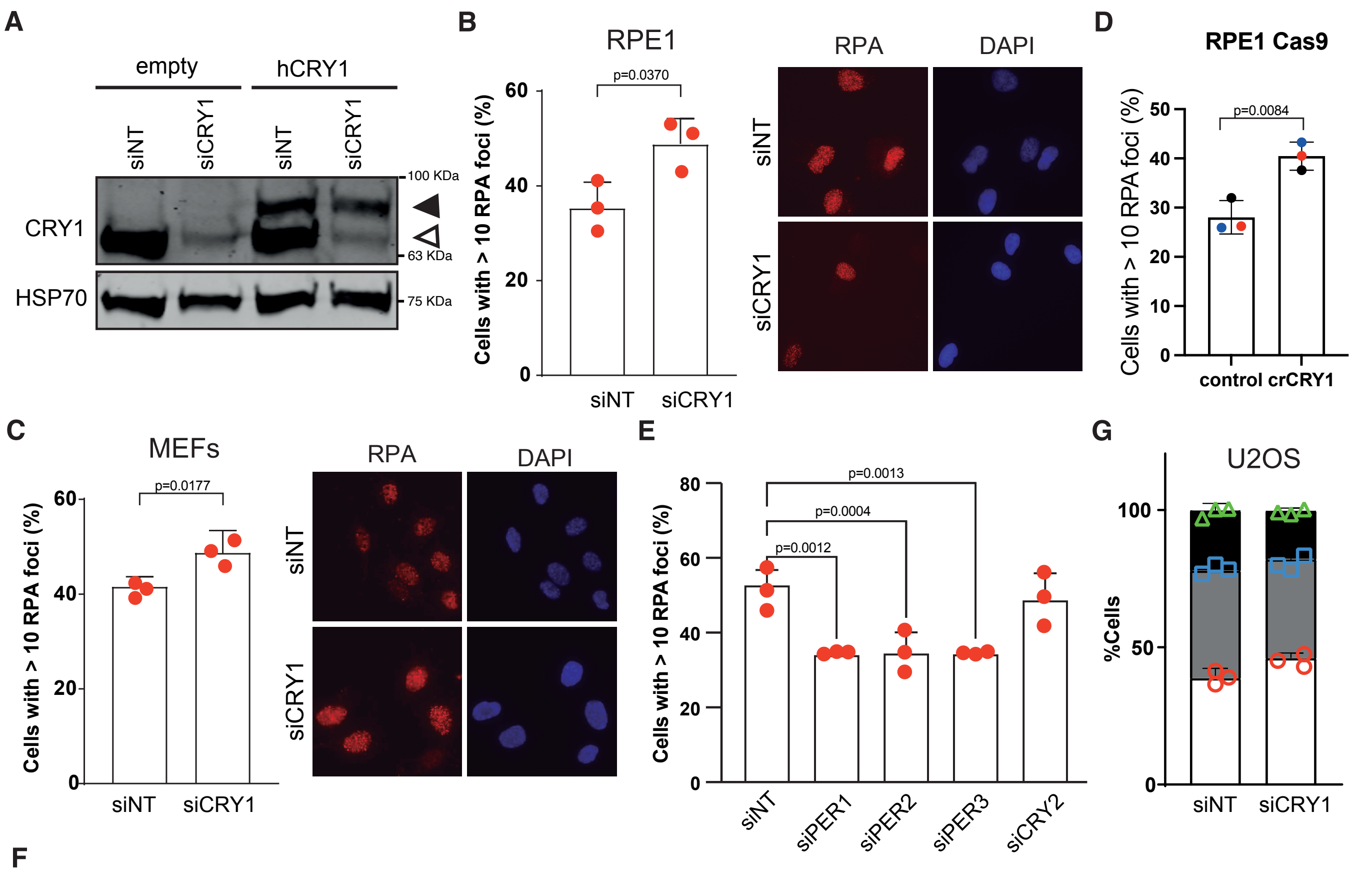


F

| | | siNT | | | | | | | siCRY1 | | | | | | |
|---|---|---|---|---|---|---|---|---|---|---|---|---|---|---|---|
| | | 0 | 4 | 8 | 12 | 16 | 20 | 24 | 0 | 4 | 8 | 12 | 16 | 20 | 24 |
| siNT | 0 | nd | ns | ns | 0.04 | ns | ns | ns | ns | ns | ns | ns | ns | ns | ns |
| | 4 | | nd | ns | ns | ns | ns | ns | ns | ns | ns | ns | ns | ns | ns |
| | 8 | | | nd | ns | ns | ns | ns | 0.03 | <0.01 | <0.01 | 0.03 | ns | 0.01 | 0.02 |
| | 12 | | | | nd | ns | ns | <0.01 | <0.01 | <0.01 | <0.01 | <0.01 | <0.01 | <0.01 | <0.01 |
| | 16 | | | | | nd | ns | ns | <0.01 | <0.01 | <0.01 | 0.03 | ns | 0.01 | 0.02 |
| | 20 | | | | | | nd | ns | ns | ns | ns | ns | ns | ns | ns |
| | 24 | | | | | | | nd | ns | ns | ns | ns | ns | ns | ns |
| siCRY1 | 0 | | | | | | | | nd | ns | ns | ns | ns | ns | ns |
| | 4 | | | | | | | | | nd | ns | ns | ns | ns | ns |
| | 8 | | | | | | | | | | nd | ns | ns | ns | ns |
| | 12 | | | | | | | | | | | nd | ns | ns | ns |
| | 16 | | | | | | | | | | | | nd | ns | ns |
| | 20 | | | | | | | | | | | | | nd | ns |
| | 24 | | | | | | | | | | | | | | nd |

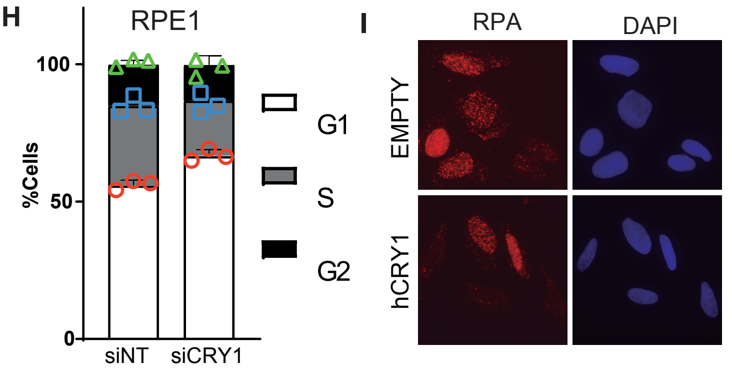


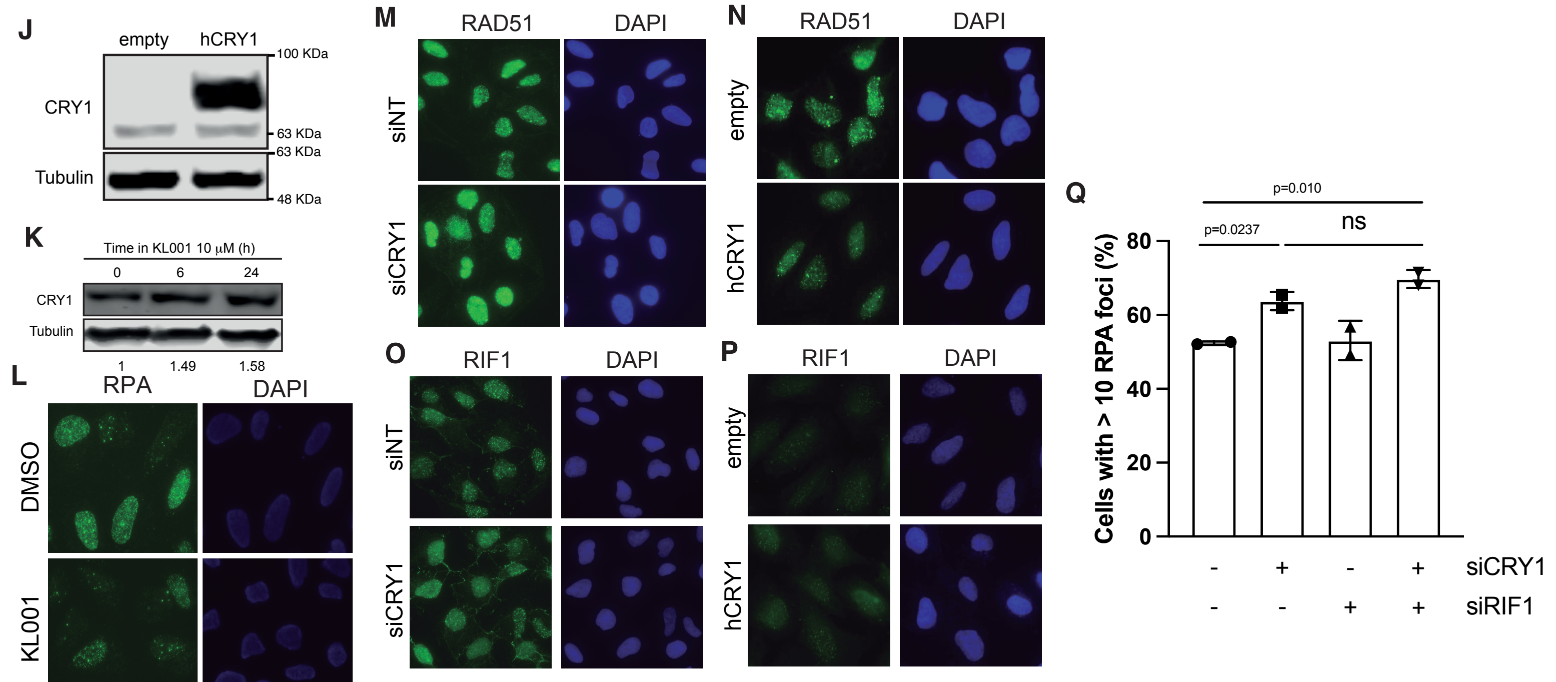


Romero-Franco et.al. Sup Figure 2

**A**

| | | siNT | | | | | | | siCtIP | | | | | | |
|---|---|---|---|---|---|---|---|---|---|---|---|---|---|---|---|
| | | 0 | 4 | 8 | 12 | 16 | 20 | 24 | 0 | 4 | 8 | 12 | 16 | 20 | 24 |
| siNT | 0 | nd | ns | ns | 0.03 | ns | ns | ns | <0.01 | 0.01 | <0.01 | <0.01 | <0.01 | <0.01 | <0.01 |
| | 4 | | nd | ns | ns | ns | ns | ns | 0.01 | ns | 0.01 | ns | 0.01 | 0.02 | 0.02 |
| | 8 | | | nd | ns | ns | ns | ns | ns | ns | ns | ns | ns | ns | ns |
| | 12 | | | | nd | ns | ns | <0.01 | ns | ns | ns | ns | ns | ns | ns |
| | 16 | | | | | nd | ns | ns | ns | ns | ns | ns | ns | ns | ns |
| | 20 | | | | | | nd | ns | <0.01 | 0.03 | <0.01 | 0.03 | <0.01 | <0.01 | <0.01 |
| | 24 | | | | | | | nd | <0.01 | <0.01 | <0.01 | <0.01 | <0.01 | <0.01 | <0.01 |
| siCtIP | 0 | | | | | | | | nd | ns | ns | ns | ns | ns | ns |
| | 4 | | | | | | | | | nd | ns | ns | ns | ns | ns |
| | 8 | | | | | | | | | | nd | ns | ns | ns | ns |
| | 12 | | | | | | | | | | | nd | ns | ns | ns |
| | 16 | | | | | | | | | | | | nd | ns | ns |
| | 20 | | | | | | | | | | | | | nd | ns |
| | 24 | | | | | | | | | | | | | | nd |

**B**

| | | shNT | | | shCTIP | | | shCCAR2 | | |
|---|---|---|---|---|---|---|---|---|---|---|
| | | 0 | 12 | 24 | 0 | 12 | 24 | 0 | 12 | 24 |
| shNT | 0 | nd | 0.0264 | ns | 0.0033 | 0.0479 | 0.0236 | ns | ns | ns |
| | 12 | | nd | 0.0142 | ns | ns | ns | 0.0481 | 0.0147 | 0.0339 |
| | 24 | | | nd | 0.0002 | 0.0034 | 0.0173 | ns | ns | ns |
| shCtIP | 0 | | | | nd | ns | ns | 0.0009 | 0.0016 | 0.0003 |
| | 12 | | | | | nd | ns | 0.0213 | 0.0387 | 0.0082 |
| | 24 | | | | | | nd | 0.0482 | 0.0137 | 0.0374 |
| shCCAR2 | 0 | | | | | | | nd | ns | ns |
| | 12 | | | | | | | | nd | ns |
| | 24 | | | | | | | | | nd |

**C**

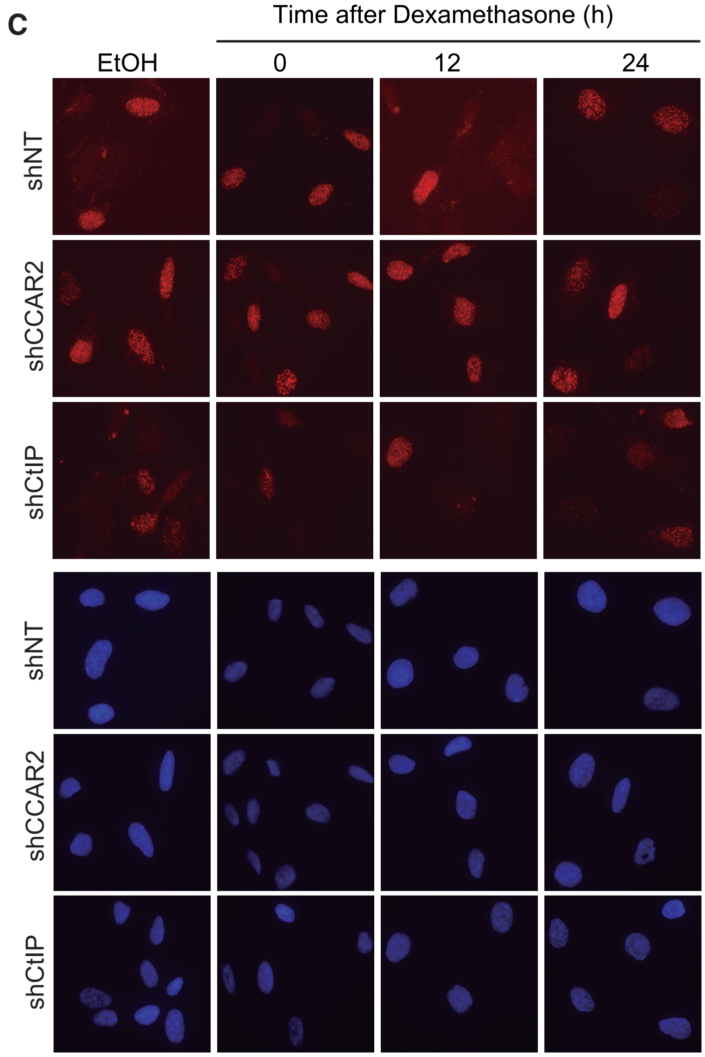


**D**

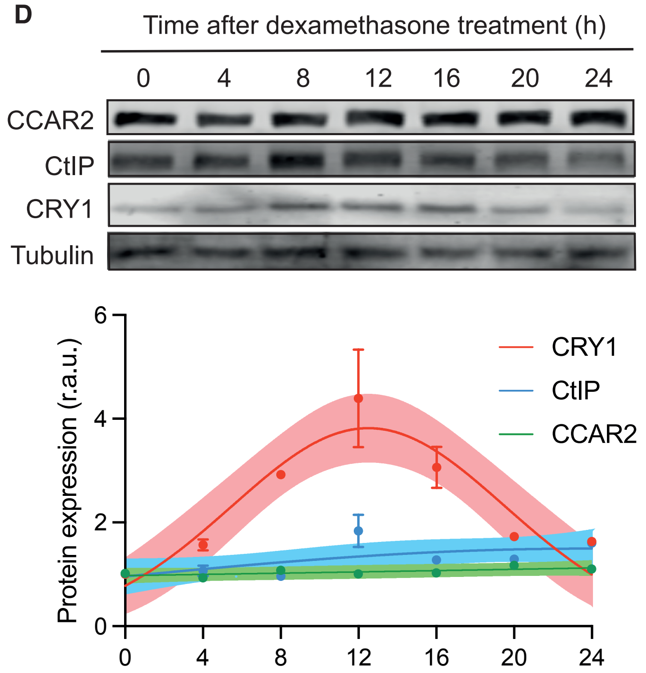


**E**

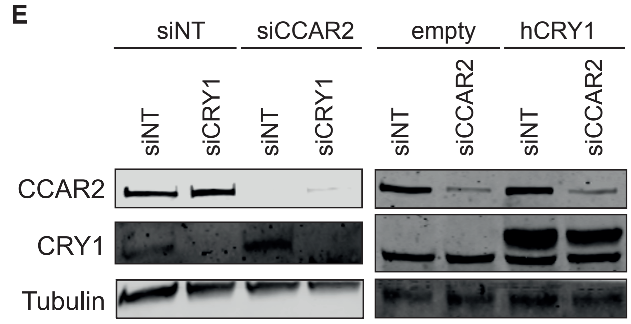


**F**

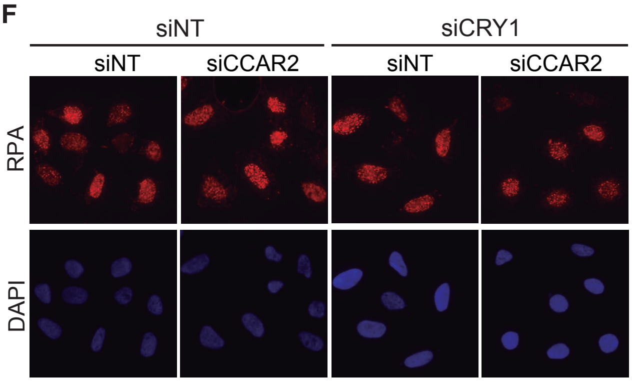


**G**

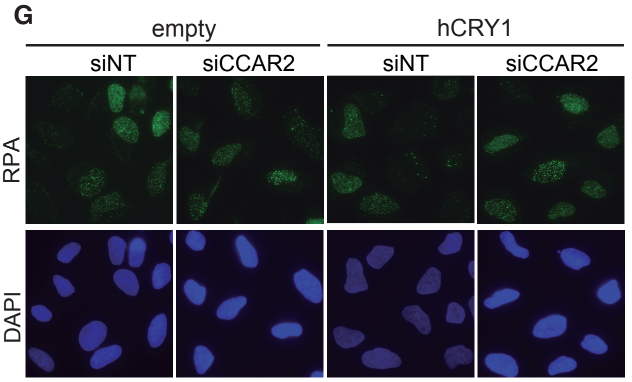


**H**

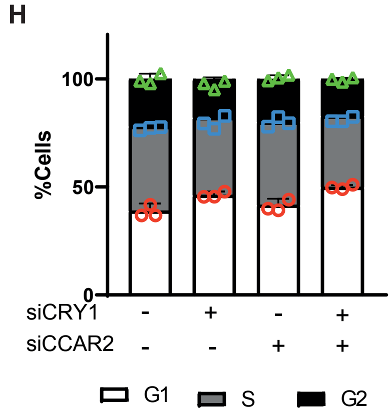


**I**

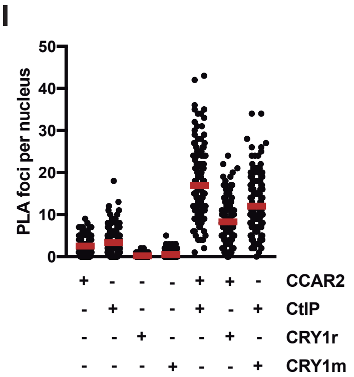


**J**

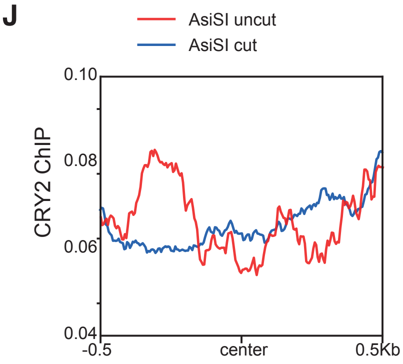


Romero-Franco et.al. Sup Figure 3

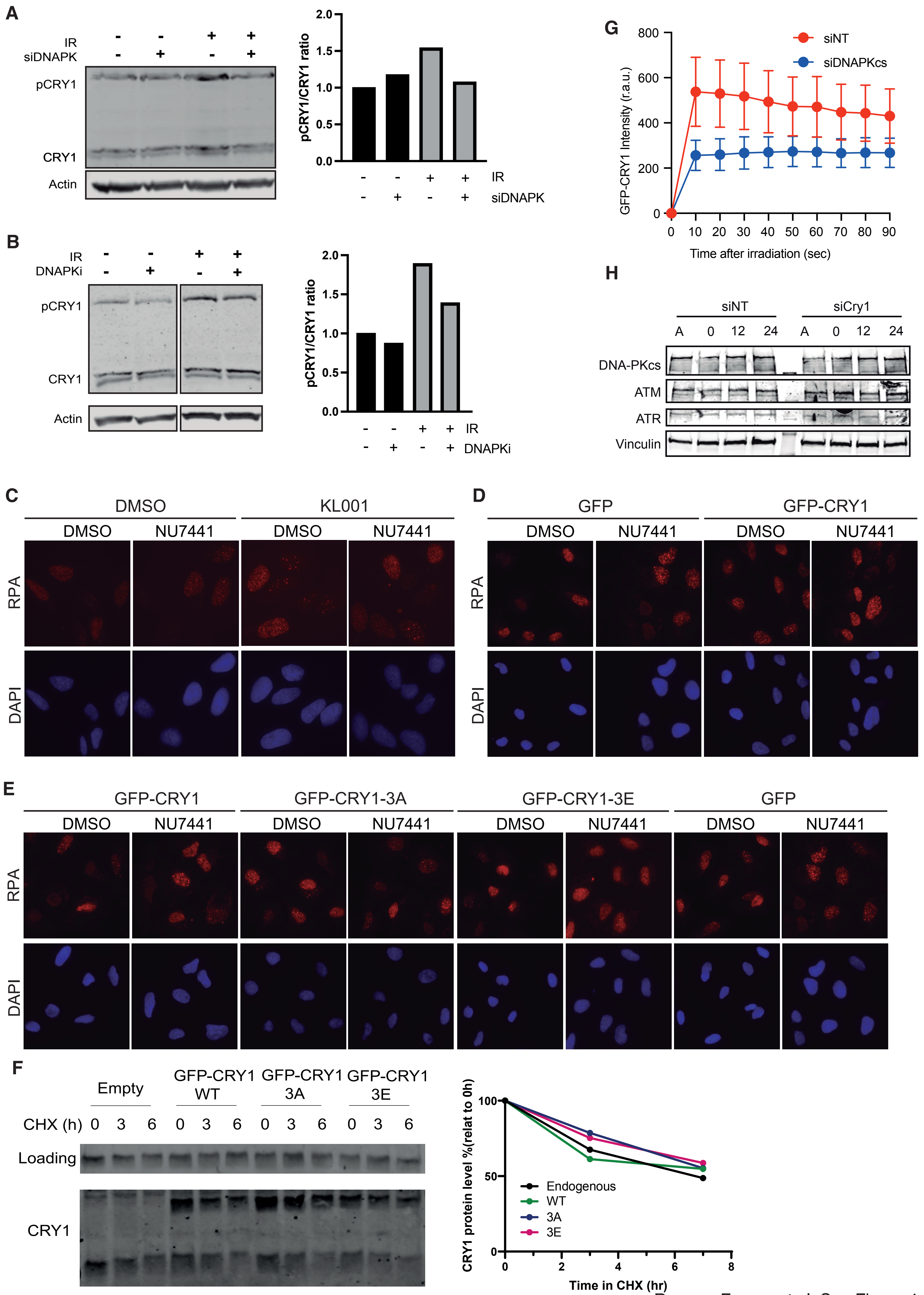

Romero-Franco et.al. Sup Figure 4

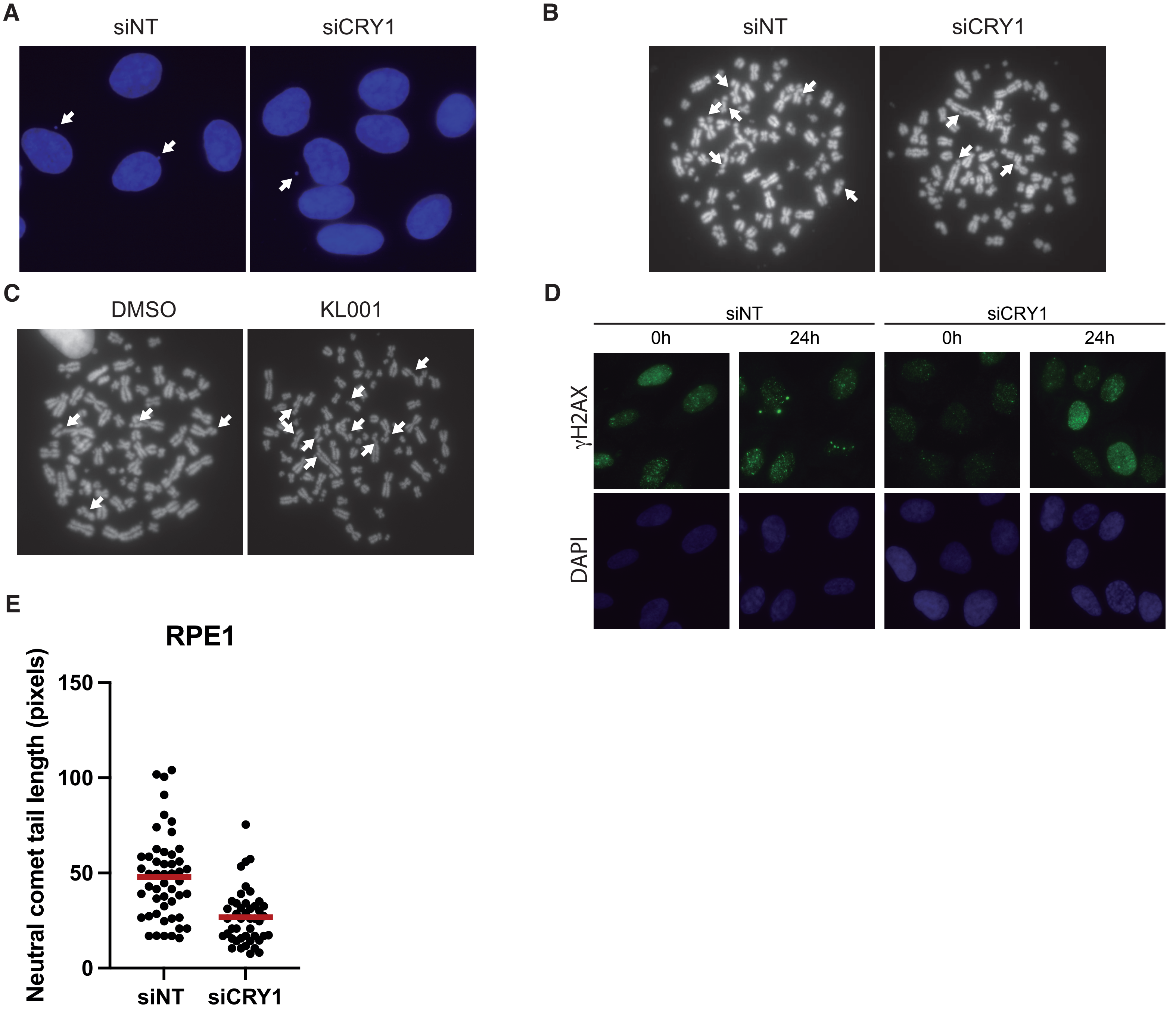
A
siNT
siCRY1
B
siNT
siCRY1
C
DMSO
KL001
D
siNT
siCRY1
0h
24h
0h
24h
γH2AX
DAPI
E
RPE1
Neutral comet tail length (pixels)
150
100
50
0
siNT
siCRY1

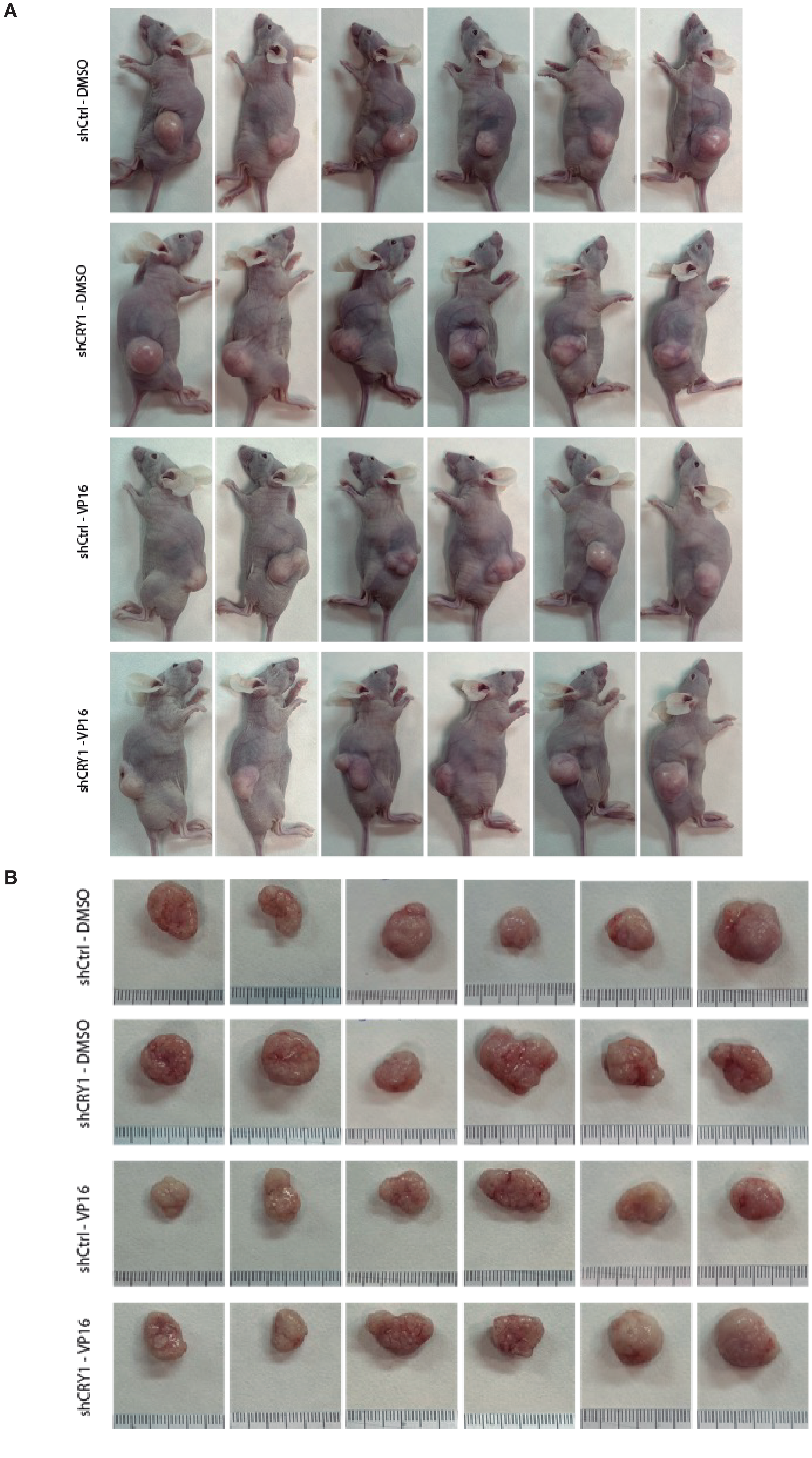


Romero-Franco et.al. Sup Figure 6
